# Structural and biochemical mechanisms of NLRP1 inhibition by DPP9

**DOI:** 10.1101/2020.08.13.250241

**Authors:** Menghang Huang, Xiaoxiao Zhang, Toh Gee Ann, Qin Gong, Jia Wang, Zhifu Han, Bin Wu, Franklin Zhong, Jijie Chai

**Affiliations:** Beijing Advanced Innovation Center for Structural Biology, Tsinghua- Peking Joint Center for Life Sciences, Center for Plant Biology, School of Life Sciences, Tsinghua University, 100084 Beijing, China; Lee Kong Chian School of Medicine, Nanyang Technological University, 11 Mandalay Road, 308232, Singapore; Skin Research Institute of Singapore (SRIS), 8A Biomedical Grove, #06-06 Immunos, 138648, Singapore; School of Biological Sciences, Nanyang Technological Unviersity; U Institute of Structural Biology, Nanyang Technological University; Max Planck Institute for Plant Breeding Research, 50829 Cologne, Germany; Institute of Biochemistry, University of Cologne, Zuelpicher Strasse 47, 50674 Cologne, Germany

## Abstract

The nucleotide-binding domain (NBD) and leucine-rich repeat (LRR)-containing receptors (NLRs) mediate innate immunity by forming inflammasomes. Activation of the NLR protein NLRP1 requires auto-cleavage within its FIIND domain^1–7^. In resting cells, the dipeptidyl peptidase DPP9 interacts with NLRP1-FIIND and together with a related enzyme DPP8, suppresses spontaneous NLRP1 activation^8,9^. The mechanisms of DPP8/9-mediated NLRP1 inhibition, however, remain elusive. Here we provide structural and biochemical evidence demonstrating that rat NLRP1 (rNLRP1) interacts with rDPP9 in a stepwise manner to form a 2:1 complex. An auto-inhibited rNLRP1 molecule first interacts with rDPP9 via its ZU5 domain. This 1:1 rNLRP1-rDPP9 complex then captures the UPA domain of a second rNLRP1 molecule via a UPA-interacting site on DPP9 and dimeric UPA-UPA interactions with the first rNLRP1. The 2:1 rNLRP1-rDPP9 complex prevents NLRP1 UPA-mediated higher order oligomerization and maintains NLRP1 in the auto-inhibited state. Structure-guided biochemical and functional assays show that both NLRP1-binding and its enzymatic activity are required for DPP9 to suppress NLRP1, supporting guard-type activation of the NLR. Together, our data reveal the mechanism of DPP9-mediated inhibition of NLRP1 and shed light on activation of the NLRP1 inflammasome.

---

In the mammalian innate immune system, detection of pathogen- or host-derived signals by NLRs induces their oligomerization, forming multiprotein complexes called inflammasomes that mediate inflammatory cell death and cytokine secretion^10,11^. NLRs generally consist of an N-terminal signaling domain, a central nucleotide-binding and oligomerization domain (NOD) and a C-terminal LRR domain. Together with a less understood sensor protein CARD8, NLRP1 (Ref. 4,7) is one of the two conserved NLR proteins harbouring an unusual domain known as FIIND (function-to-find domain)^12^ (Fig. 1a). The FIIND has intrinsic proteolytic activity and undergoes auto-proteolysis between two subdomains: ZU5 (found in ZO-1 and UNC5) and UPA (conserved in UNC5, PIDD, and Ankyrin). Although the cleaved ZU5- and UPA-containing fragments remain non-covalently associated, the auto-cleavage is a pre-requisite for NLRP1 (Ref. 4,7) and CARD8 activation ^13^. *Bacillus anthracis* lethal factor (LF) is the best characterized pathogen-derived trigger for rodent NLRP1 (Ref. 5,14). LF cleaves mNlrp1b close to its N-terminus and induces proteasomal degradation of the entire N-terminal NOD-LRR-ZU5 fragment via the N-end rule pathway^1–3^. This liberates the active UPA-CARD fragment that rapidly oligomerizes to engage downstream inflammasome effectors such as ASC and pro-caspase-1 (Ref. 1,3). This unique mechanism involving ‘functional degradation’ is conserved between human and rodent NLRP1 homologues and also governs the activation of human CARD8 (Ref. 11).

**Fig. 1.**
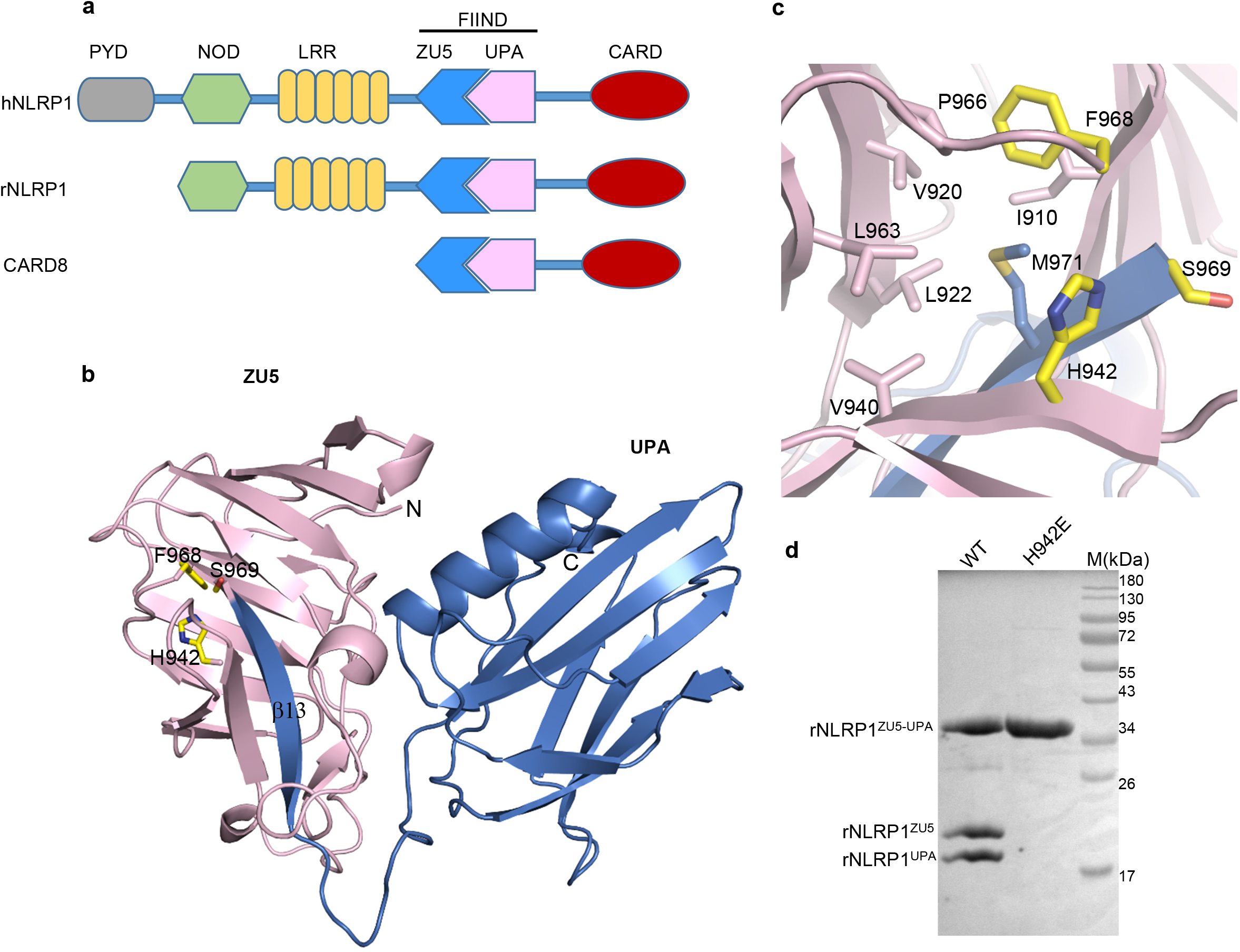
Crystal structure of the FIIND domain of rNLRP1. **a**, Schematic diagram of domain structures of human NLRP1, rat NLRP1 and CARD8. PYD: pyrin domain; CARD: caspase activation and recruitment domain. **b**, Crystal structure of the FIIND domain of rNLRP1. The ZU5 and UPA domains are shown in pink and blue, respectively. The catalytic residues of the FIIND are labeled and shown in stick mode. **c**, A close-up view of the catalytic site of the FIIND. **d**, Mutation of the catalytic residue His942 abolishes autoproteolysis of the FIIND domain of rNLRP1. Wild type and the His942 mutant rNLRP1^FIIND^ proteins were purified from insect cells and visualized by SDS-PAGE (polyacrylamide gel electrophoresis) followed by Coomassie Blue staining.

Dipeptidyl peptidases 8 and 9 (DPP8/9) are highly related intracellular prolyl peptidases and implicated in immunoregulation^15^. They act as endogenous inhibitors of the NLRP1 inflammasome in human^8,9,13^ and rodent^6,8,13,16,17^. Notably, the FIIND of hNLRP1 is necessary and sufficient for interaction with hDPP9 (Ref. 9). Furthermore, inhibitors of class IV DPPs, such as Valine-boro-Proline (VbP), or CRISPR/Cas9-mediated DPP8/9 knockout specifically activate NLRP1 and/or CARD8 (Ref. 6,8,9,13). Like LF, VbP also induces proteasome-mediated degradation of mNlrp1b N-terminal fragment. However, unlike LF, this does not require N-degron recognition. Hence VbP and LF trigger NLRP1 activation in two distinct, independent pathways. Overall, the mechanisms by which DPP9 prevents spontaneous inflammasome activation are not well understood.

Here we elucidate the structural mechanisms by which DPP9 suppresses NLRP1 activation via the FIIND domain. A cryo-EM structure of the complex containing full-length rNLRP1 and full-length rDPP9 determined at 3.18 Å showed that rNLRP1 and rDPP9 formed a complex with a stoichiometry of 2:1. Mutations at a single rNLRP1-rDPP9 interface completely disrupted the complex, suggesting that the complex is formed by stepwise, sequential binding of two rNLRP1 molecules with distinct conformations and modes of interaction. Furthermore, the 2:1 rDPP9-rNLRP1 complex prevents higher order UPA oligomerization, which is a pre-requisite for NLRP1 activation. More importantly, biochemical and functional data showed that NLRP1 binding and peptidase activity of DPP9 are both indispensable in preventing spontaneous NLRP1 inflammasome activation in cells. These results resolve the structural mechanisms by which NLRP1-FIIND orchestrates the switch between auto-inhibition and activation via DPP9 association. We propose that the 2:1 rNLRP1-rDPP9 complex is a bona fide innate immune surveillance complex in which NLRP1 acts as a guard for DPP9 function.

## Crystal structure of auto-inhibited FIIND

Previously it was shown that the ZU5 subdomain is sufficient to block the release of the active UPA-CARD fragment of NLRP1 (Ref. 3,9). To elucidate the molecular basis of ZU5 domain-mediated NLRP1 auto-inhibition, we crystallized the FIIND domain from rNLRP1 and solved its structure using molecular replacement (Extended Data Table 1). Electron density unambiguously shows that the purified FIIND is auto-cleaved at the predicted position between Phe968 and Ser969 (Extended Data Fig. 1a). This is further confirmed by SDS PAGE of the crystals (Extended Data Fig. 1b). Positioning of ZU5 and UPA resembles that of the two structural domains in the auto-inhibited UNC5b (Extended Data Fig. 1c)^18^. Inter-domain interaction largely comes from the first β-strand (β13) of UPA, which forms two anti-β sheets with ZU5 (Fig. 1b). This accounts for the tight association of the two cleaved fragments of NLRP1 and why the ZU5 domain can block the release of the active UPA-CARD fragment.

The catalysis-essential ‘FS’ motif^19^ is conserved in NLRP1 homologues and CARD8 (Extended Data Fig. 1d). In the structure, rNLRP1^F968^ from this motif points into a pocket created by the side-chains of several hydrophobic residues (Fig. 1c), suggesting that this residue provides a structural anchor to stabilize the local conformation. A similar role has been described for the corresponding phenylalanine in the FIIND-containing proteins Nup98 (Ref. 20) and PIDD^21^. Intriguingly, rNLRP1^His942^, highly conserved in NLRP1 and CARD8 (Extended Data Fig. 1d), is located adjacent to the serine residue (rNLRP1^S969^) from the ‘FS’ motif (Fig. 1b). Mutation of this histidine residue resulted in a loss of rNLRP1 autocleavage when expressed in insect cells (Fig. 1d). A similar effect was also observed for mutations of corresponding CARD8^H270^ or hNLRP1^H1186^ (Ref. 19). These data suggest that rNLRP1^H942^ is a catalytic residue, which likely functions to activate the catalytic rNLRP1^S969^ for auto-proteolysis.

## Architecture of the 2:1 rNLRP1-rDPP9 complex

To probe the mechanism of NLRP1 inhibition by DPP9, we co-expressed full-length rNLRP1 and rDPP9 in insect cells and purified them using affinity chromatography. Supporting previous studies^3,8,9^, gel filtration analysis indicated that the two proteins formed a stable complex (Extended Data Fig. 2a). The rNLRP1-rDPP9 complex purified from gel filtration was then used for structural analysis with cryo-EM. 2D averages showed a dimeric DPP9 but only one subunit bound NLRP1 in most of the particles (Extended Data Fig. 2c), suggesting that some rNLRP1-rDPP9 complex had dissociated during cryo-EM sample preparation. We used the particles with one rDPP9 subunit bound by rNLRP1 for further cryo-EM analysis. After three-dimensional (3D) classification, a subset of 182,116 particles were used for reconstruction, generating a map with a global resolution of 3.18 Å (Fig. 2a, Extended Data Fig. 2c-f and Extended Data Table 2) as determined with a gold-standard Fourier shell correlation.

**Fig. 2.**
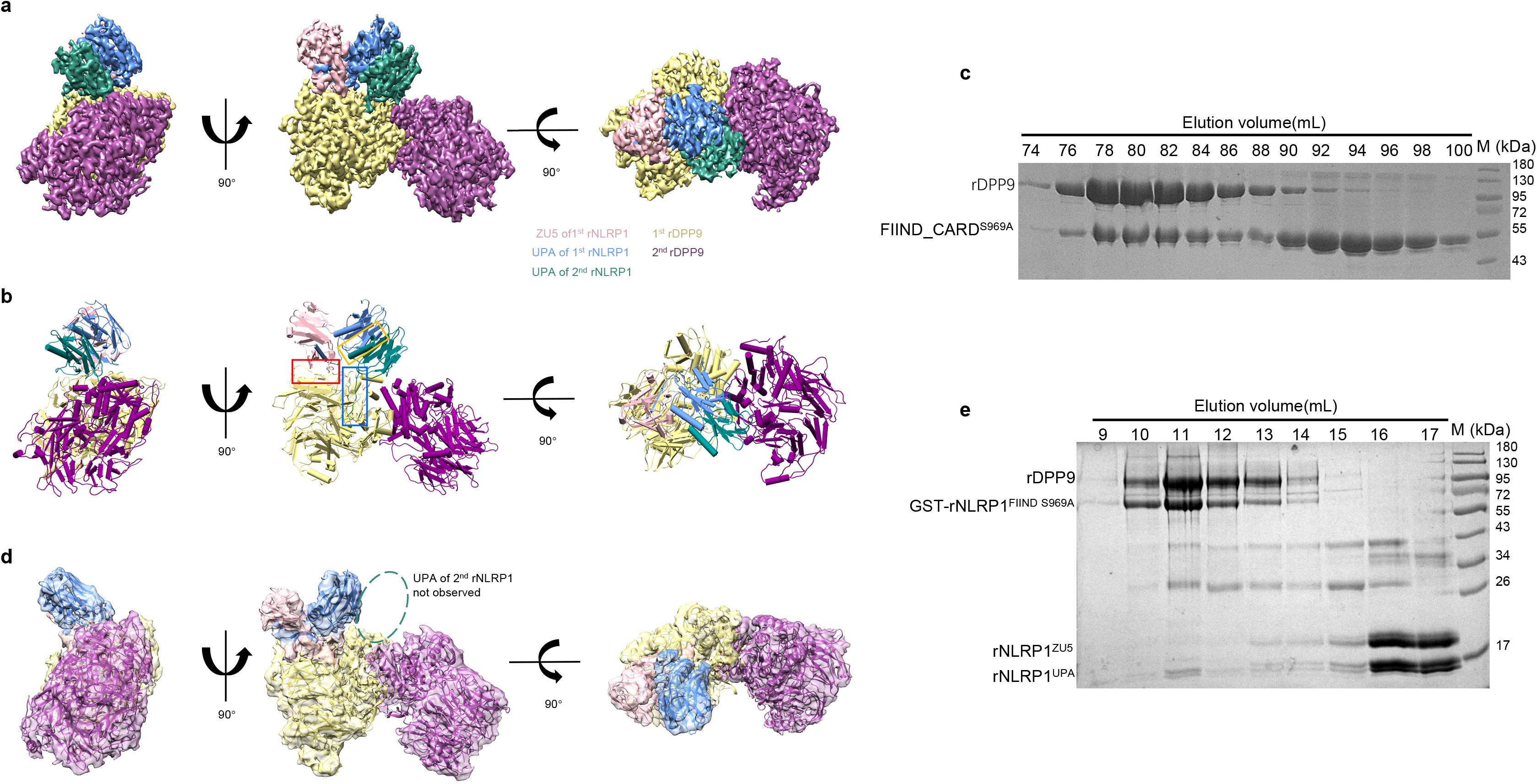
Architecture of the 2:1 rNLRP1-rDPP9 complex. **a**, The final cryo-EM density of the rNLRP1-rDPP9 complex at 3.18 Å shown in three orientations. Color codes for domain structures are indicated. **b**, Model of the rNLRP1-rDPP9 complex shown in three orientations. The three distinct interfaces are shown in colored boxes. Red, ZU5 binding site; Blue, UPA binding site; Yellow, UPA-UPA binding site. **c**, Autoproteolysis is dispensable for rNLRP1^FIIND^ interaction with rDPP9. The mutant S969A of N-terminally His-tagged rNLRP1^FIIND-CARD^ and non-tagged rDPP9 were co-expressed in insect cells and purified using affinity chromatography and gel filtration. Shown in the figure are fractions of the two proteins after gel filtration visualized by SDS-PAGE followed by Coomassie Blue staining. **d**, 3D reconstruction of the rNLRP1^FIIND S969A^-rDPP9 complex shown in three orientations. Highlighted within the open ellipse is the vacant UPA-binding site. **e**, The ZU5 subdomain is displaced following interaction of the second rNLRP1 with rDPP9. Pre-formed rNLRP1^FIIND S969A^–rDPP9 complex was incubated with fully auto processed rNLRP1^FIIND^ and analyzed with gel filtration. Protein fractions from the gel filtration assay were visualized by SDS-PAGE followed by Coomassie-blue staining.

rDPP9 forms a homodimer (Fig. 2b) that is nearly identical with the dimeric hDPP9 (Ref. 22). Several flexible loop regions were not observed in the reported free hDPP9 structure, but their equivalents are well defined in the rNLRP1-bound rDPP9 subunit, suggesting that interaction with rNLRP1 stabilizes these loop regions (Extended Data Fig 4a). A similar observation was previously made with substrate-bound hDPP9 (Ref. 22).

Unexpectedly, our cryo-EM structure revealed that the liganded rDPP9 subunit is bound by two rNLRP1 molecules (Fig. 2b), hereby termed the 2:1 rNLRP1-rDPP9 complex. Furthermore, the two rNLRP1 molecules adopt distinct conformations and modes of interaction with rDPP9. The first bound rNLRP1 molecule contains the complete FIIND domain in a conformation nearly identical to that of the free FIIND crystal structure (Extended Data Fig. 4b). Except for the FIIND, none of the remaining domains of this rNLRP1 molecule is well defined in the cryo-EM density, suggesting that they are flexible relative to the FIIND. In the 2^nd^ bound rNLRP1 molecule, only the UPA domain within the FIIND is well defined, suggesting that the N-terminal domains including ZU5 have dissociated from the UPA when bound in this position (see further discussion below). In support of this, the post-cleavage N-terminal NOD-LRR-ZU5 fragment was present in sub-stochiometric ratios relative to the C-terminal UPA-CARD fragment in the rNLRP1-rDPP9 complex (Extended Data Fig.2a).

Three key interaction surfaces are responsible for the formation of the unexpected 2:1 rNLRP1-rDPP9 complex (Extended Data Fig 8d). 1). The first rNLRP1 molecule interacts with rDPP9 via its ZU5 domain, which binds to one lateral side of the β-propeller domain of rDPP9 (herein termed ZU5-binding site on rDPP9 (Fig. 2b, red box). 2) The UPA domain of the second bound rNLRP1 molecule interacts with rDPP9 by capping the opening of the rDPP9 active site and extending an N-terminal loop deep into the rDPP9 substrate binding channel. Hence, the second rNLRP1 binding surface on rDPP9 (termed herein UPA-binding site) (Fig. 2b, blue box) overlaps extensively with the rDPP9 catalytic groove. 3) The UPA domains of the two bound rNLRP1 molecules establish extensive interactions with each other via a new homodimerization surface (termed UPA dimerization site) (Fig. 2b, yellow box). Remarkably, the loop segment of the UPA domain used by the second rNLRP1 molecule to interact with the rDPP9 UPA-binding site normally forms a β-sheet with ZU5 in the free FIIND (Fig. 1b). This suggests that rDPP9 binding disrupts the intermolecular interactions of the two cleaved FIIND fragments (i.e. ZU5 and UPA) when rNLRP1 is bound in the second position.

## Distinct binding modes of the two bound rNLRP1 molecules

Previous data indicated that protease activity is not required for DPP9 binding to NLRP1 and CARD8 (Ref. 8,9). Indeed, gel filtration showed that the catalytically inactive rDPP9^S729A^ mutant formed a stable complex with the rNLRP1 FIIND-CARD fragment (Extended Data Fig. 5a). In addition, in the first bound position (Fig. 2b), the rNLRP1 FIIND domain adopts a near identical conformation with the free rNLRP1 FIIND molecule in the crystal structure (Extended Data Fig. 4b). These results predict that a non-autocleavable rNLRP1, which does not support rNLRP1 activation in cells^7,19^, can still interact with rDPP9 via the ZU5 domain. As expected, the FIIND-CARD cleavage mutant S969A (FIIND-CARD^S969A^) retained the activity of interacting with rDPP9 though it underwent no auto-proteolysis (Fig. 2c). These results are consistent with a previous study of CARD8 FIIND ^8,13^, which interacts with hDPP9 regardless of auto-cleavage.

In contrast to the first bound position, the second rNLRP1 molecule in the 2:1 complex employs an N-terminal loop segment in its UPA to interact with rDPP9, which is mutually exclusive with its normal role (Fig. 2b) in forming an intermolecular β-sheet with its own ZU5 domain (Fig. 1b). This indicates that in direct contrast to the first position, rNLRP1 FIIND must adopt a different conformation from the free form in order to dock into the second position. To test this hypothesis biochemically, we performed cryo-EM analysis for rDPP9 in complex with FIIND-CARD^S969A^, which cannot undergo autoproteolysis and therefore mimics the auto-inhibited, free FIIND conformation (Extended Data Fig.3). The mutant FIIND-CARD^S969A^ formed a strong 1:1 complex with rDPP9. The structure of the mutant complex is nearly identical to that of the wild-type complex encompassing the first rNLRP1 molecule, except no clear density was found in the second position (Fig. 2d). These results indicate that FIIND-CARD^S969A^ is competent to occupy the first binding position using the ZU5 domain, but cannot interact with the UPA-binding site in the second position. They may also explain why autoproteolysis-deficient NLRP1 mutants still retain but have reduced DPP9-binding activity^8,9^. These results also establish that rNLRP1 must undergo an auto-proteolysis dependent conformation shift in order to fit into the second position in the 2:1 complex.

The interactions between the first rNLRP1 molecule and the ZU5-interacting site on rDPP9 consist of extensive contacts between β10, α1-β6 loop of rNLRP1 FIIND ZU5 and the β-propeller domain of rDPP9 (Fig. 3a,b). FIIND^β10^ makes three main-chain hydrogen bonds with the N-terminal portion of a long loop of rDPP9, forming an anti-β-sheet like structure (Fig. 3b). The long rDPP9 loop also makes hydrophobic contacts with the middle part of the α-β6 loop of FIIND. Notably, the amino acids of this ZU5-interacting site are missing in DPP4 but highly conserved in DPP8 (Extended Data Fig. 4c), explaining why DPP8 but not DPP4 interacts with and inhibits Nlrp1b^8,23^. In addition, the short α-helix in the FIIND α-β6 loop packs against other two loops of rDPP9. One of these two loops establishes additional interactions with the inter-domain loop region of FIIND (Fig. 3b).

**Fig. 3.**
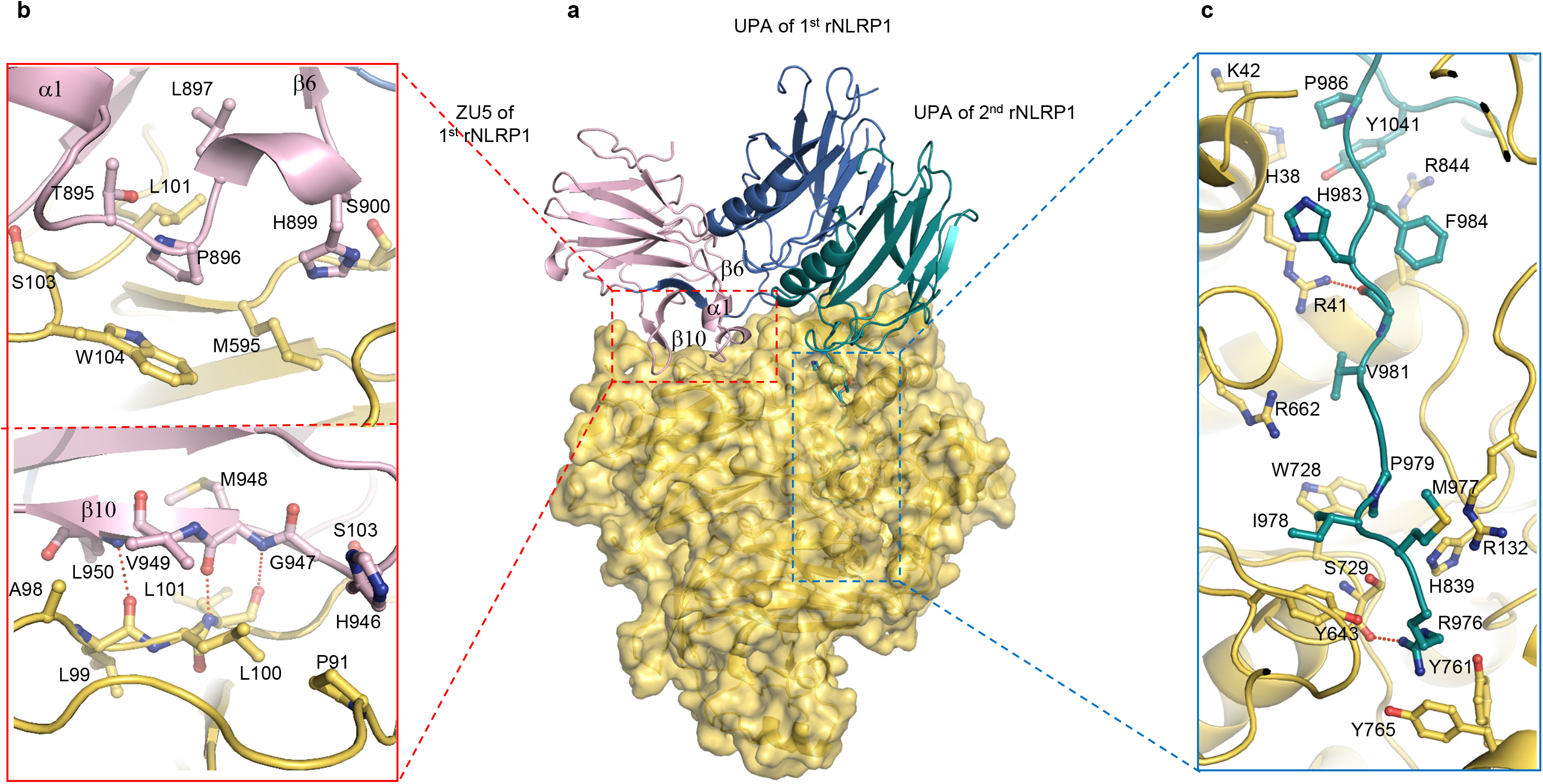
Molecular composition of the two distinct rNLRP1-rDPP9 binding sites. **a**, Structure of the rNLRP1-rDPP9 complex. The critical regions involved in rDPP9 binding in the ZU5 domain of the 1st rNLRP1 molecule are labelled, and the interacting regions between domains are highlighted with open frames. **b**, Detailed interactions between ZU5 and rDPP9 at the ZU5-binding site for the red-framed region. Hydrogen bonding interaction are indicated by red dashed lines. **c**, Detailed interactions between UPA and rDPP9 at the UPA-binding site for the blue-framed region.

Two possible explanations for why the ZU5 domain could not be observed in the 2nd position (Fig. 2b): 1) ZU5 becomes flexible after UPA-rDPP9 interaction, or 2) ZU5 becomes dissociated after rNLRP1 docks into the second position. To differentiate these two possibilities, we tested how an auto-cleaved FIIND-CARD fragment fits into the second position with a preformed 1:1 FIIND-CARD^S969A^-rDPP9 complex. Gel filtration showed that the auto-cleaved FIIND-CARD indeed complexed with the preformed 1:1 FIIND-CARD^S969A^-rDPP9 complex. In the resulting complex, rNLRP1-ZU5 and UPA became substoichiometric (Fig. 2e) with substantially less ZU5 than UPA-CARD indicating that the ZU5 domain becomes dissociated from the UPA after rNLRP1 docks into the second position.

Another notable feature of the second bound rNLRP1 is that the substrate binding groove of rDPP9 is completely blocked by the N-terminally extended loop of the UPA (Fig. 3a, c). The extreme N-terminal 7 residues of the UPA loop are non-structured, but they are not cleaved by DPP9 itself, as evidenced by N-terminal sequencing (Extended Data Fig. 5c), in agreement with previous cellular data on hNLRP1 (Ref. 8,9,24).

## Two rNLRP1 molecules bind to rDPP9 sequentially

As rNLRP1-FIIND^S969A^ can form a 1:1 complex with rNLRP1 with a vacant second position (Fig. 2d), we asked if the reverse is true, i.e. if the first binding position is required for the second rNLRP1 to bind rDPP9. Point mutations within the interface between the 1st rNLRP1 and rDPP9 (rNLRP1^S900^ and rDPP9^L101^) completely abrogated the interaction between rNLRP1 and rDPP9 (Fig. 4a and Extended Data Fig. 6, Fig. 4b). Similarly, a mutation of the corresponding residue of rDPP9^L101^ in hDPP9 (L131) also abrogated the interaction with hDPP9 altogether in 293T cells (Fig. 4c). Therefore, rNLRP1 cannot interact with rDPP9 solely in the second binding position. In other words, the first rNLRP1 molecule must be bound in the 1st position via the ZU5 domain before the second rNLRP1 molecule can dock into the second position via the UPA domain.

**Fig. 4.**
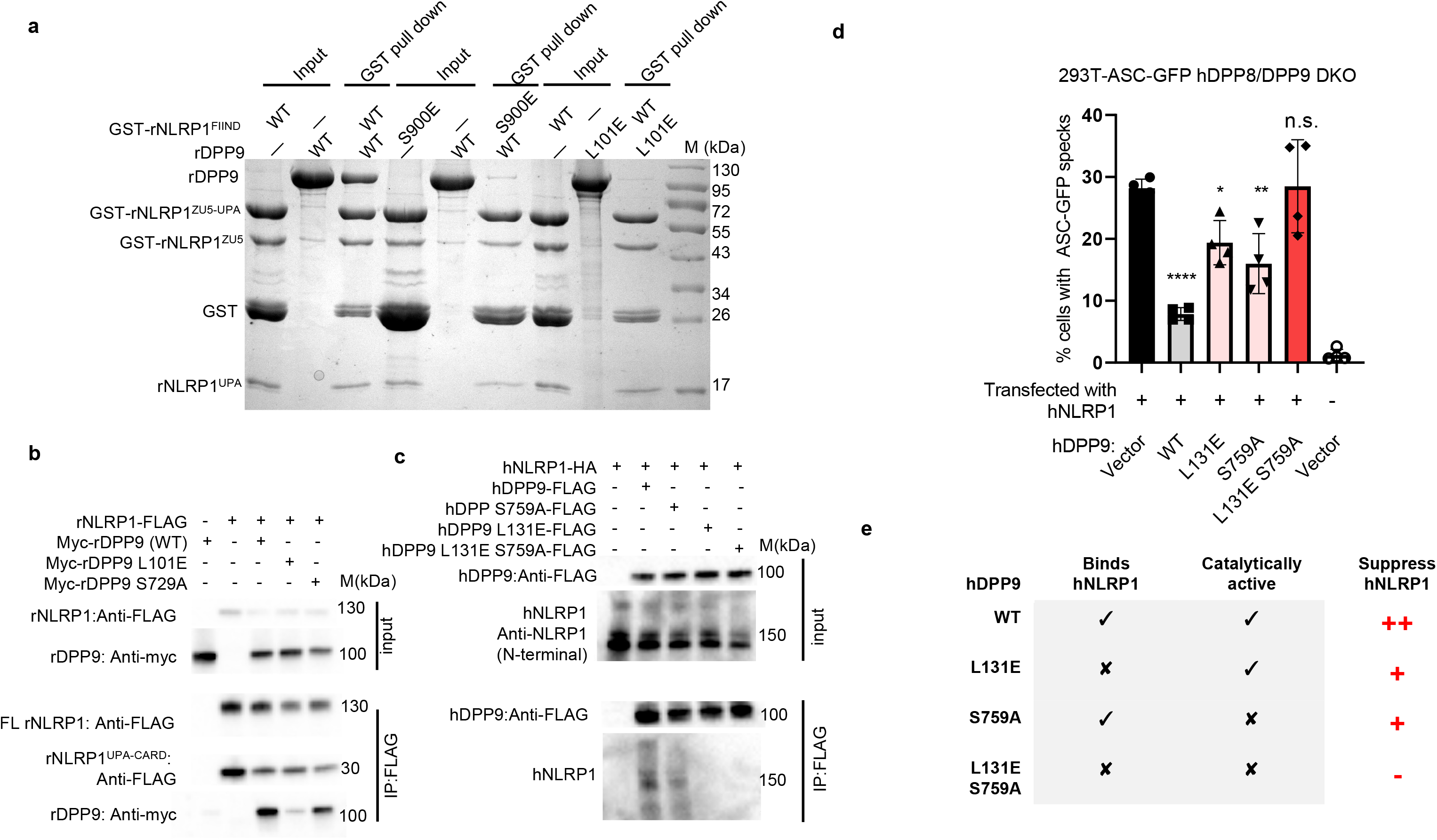
Both protease activity and FIIND binding are important for DPP9 inhibition of NLRP1. **a**, N-terminally GST-fused rNLRP1^FIIND^ was used to pull down non-tagged rDPP9 in vitro. The proteins were purified using glutathione sepharose 4B (GS4B) and proteins bound to the resin were detected by SDS-PAGE followed by Coomassie-blue staining. **b**, Lysates from 293T cells transfected with indicated FLAG-tagged rNLRP1 and myc-tagged rDPP9 were subjected to anti-FLAG IP. **c**, The assay was performed as described in (**b**) using FLAG-hDPP9 and HA-hNLRP1. **d**, 293T-ASC-GFP DPP8/9-DKO cells were transfected with constructs indicated. Y-axis represents the percentage of cells with ASC-GFP specks. Bar graphs represent data from biological triplicates. **e**, A summary of catalytic activity, hNLRP1 FIIND binding and hNLRP1 suppression of WT hDPP9, hDPP9^L131E^ and hNLRP1^S759A^.

## Role of the DPP9 catalytic activity and DPP9-FIIND interaction in NLRP1 activation

We next took advantage of the mutation hNLRP1^L131E^, which abrogates hNLRP1-hDPP9 binding, to determine whether NLRP1 binding is functionally important for DPP9 inhibition of NLRP1. To this end, we imaged ASC-GFP-expressing hDPP8/9 knockout 293T cells co-expressing hNLRP1 and hDPP9 (Ref.9). As anticipated, transfection of hDPP9 inhibited NLRP1-dependent formation of ASC-GFP specks, confirming that wild-type hDPP9 prevents spontaneous hNLRP1 activation (Fig. 4d and Extended Data Fig. 8a). The inhibition was partially but significantly retained in hDPP9^L131E^ or the catalytic mutant hNLRP1^S759A^, indicating that both catalytic activity and FIIND binding are important for hDPP9-mediated inhibition of hNLRP1 (Fig. 4d and Extended Data Fig. 8a). The effect was not caused by alteration of protease activity of the hDPP9^L131E^ mutant (Extended Data Fig. 5b). In further support of this conclusion, simultaneous mutations of these two residues completely eliminated the inhibitory effect of hDPP9 on formation of ASC specks (Fig. 4d, e). These results are broadly consistent with previous results demonstrating that hDPP9^S759A^ failed to fully rescue hNLRP1 activation in hDPP9-deficient 293T cells^8^.

DPP8/9 inhibitors like VbP have been shown to bind to the active site of hDPP9, which overlaps with the UPA-binding site. Thus, VbP might compete with NLRP1-UPA-CARD for interaction with rDPP9. However, pull-down assays showed that VbP had little effect on rNLRP1^FIIND^ interaction with rDPP9 (Extended Data Fig. 7a). This is consistent with data from mNlrp11b and CARD8 but in contrast with those from hNLRP1 (Ref. 8,9). The precise reason for this remains unclear, but it might be that the corresponding segment of hCARD8 is not involved in interaction with hDPP9 due to variations in the sequence around the N-terminal segment (Extended Data Fig. 1d). The autoinflammatory disease-associated mutation hNLRP1^P1214R^ may perturb interaction of the hNLRP1 interaction with hDPP9 at the UPA-binding site^9^.

## Structural basis for ZU5-mediated inhibition of UPA-CARD oligomerization

Finally, we examined the third critical interaction interface within the 2:1 rNLRP1-rDPP9 complex: the homodimeric UPA-UPA interface between the first and second rNLRP1 molecules. This interaction is mainly mediated by the long loop C-terminal to β7 and β8 from one side of UPA (Fig. 5a). In a recent study, we reported that hNLRP1-UPA-CARD, which represents the active fragment of NLRP1, forms filament-like higher molecular weight oligomers^3^. This process proceeds via a two-step mechanism with the UPA forming a ring-like oligomer, which then brings the CARDs in close proximity for efficient filament-like polymerization. We hypothesize that the same UPA-UPA interaction surface as observed in the 2:1 NLRP1-DPP9 complex is utilized in the higher order UPA-CARD filament. Indeed, negative staining EM showed that the wild-type UPA-CARD of hNLRP1 formed filamentous structures^25,26^ (Fig. 5b), but not the UPA-UPA interface mutants, UPA-CARD^P1278E^ and UPA-CARD^L1281E^ (Fig. 5b).

**Fig. 5.**
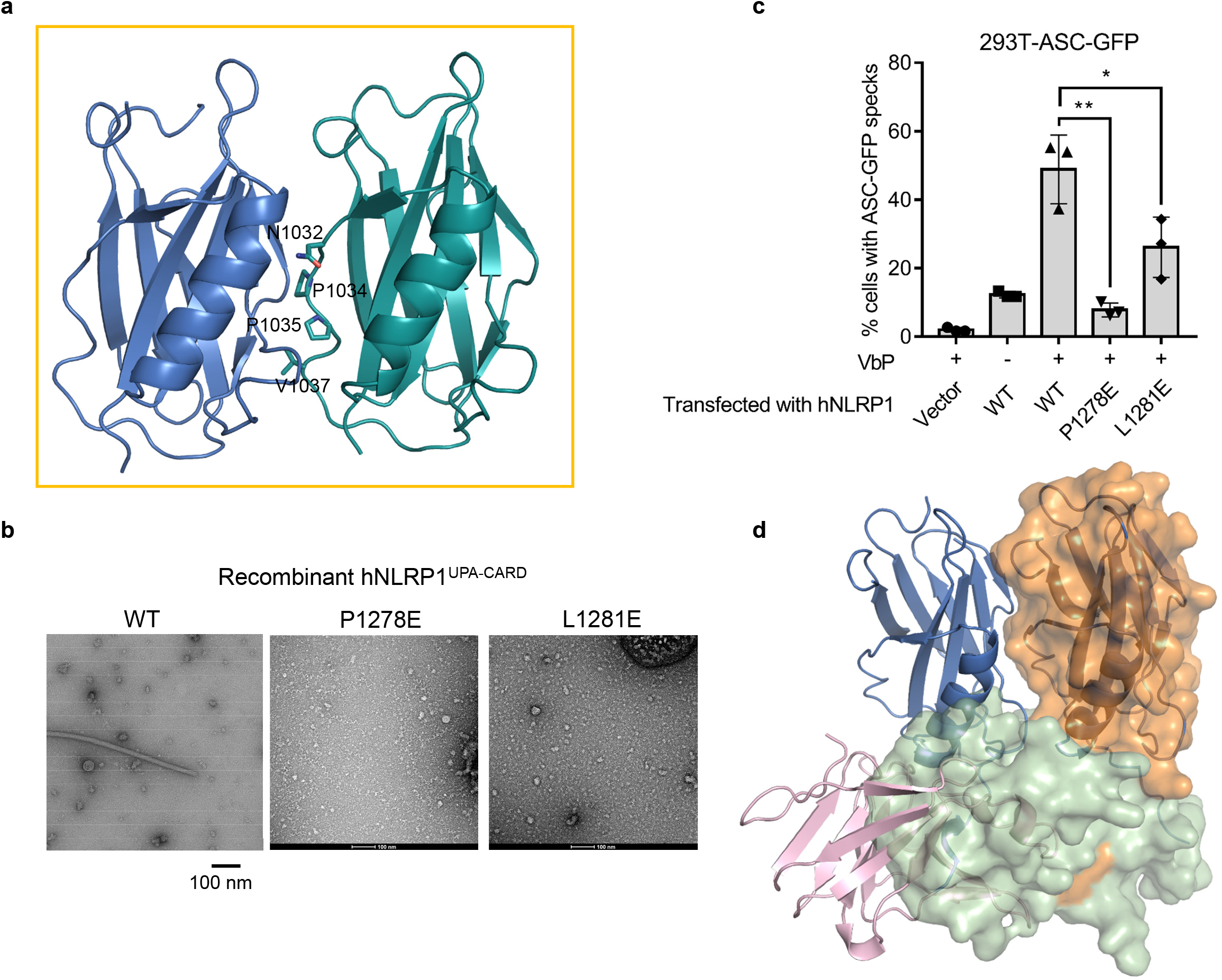
Auto-inhibition mechanism of the FIIND domain. **a**, Cartoon representation of the UPA dimer in the structure of the 2:1 rNLRP1-rDPP9 complex. Key residues mediating formation of the dimeric UPA are shown in stick. **b**, Negative staining electron microscopy (EM) analysis of human WT FIIND-CARD (left), FIIND-CARD^P1278E^ (middle) FIIND-CARD^L1281E^ (right). **c**, Impact of hNLRP1^P1278E^ and hNLRP1^L1281E^ on VbP-induced activation of hNLRP1 in 293T cells. 293T-ASC-GFP cells were transfected with empty vector or hNLRP1 plasmids indicated and treated with 5 μM VbP for 24 h. **d**, Alignment of the UPA homodimer (in cartoon) from 2:1 rNLRP1-rDPP9 complex with the crystal structure of rNLRP1^FIIND^ (in transparent surface). The UPA monomer at on the right side (from the 2^nd^ rNLRP1 in the 2:1 complex) was used as the template for the alignment.

A closer examination revealed that residues Asn1032, Pro1034, Pro1035 and Val1037, located at the center of the UPA-UPA dimer surface, are highly conserved among NLRP1 proteins from different species and CARD8 (Extended Data Fig.1d). Interestingly, an hNLRP1 splice isoform, hNLRP1-2 that lacks this loop region displayed impaired cell-killing activity when ectopically expressed in MCF7 cells^27^, supporting an essential role of this loop in hNLRP1-mediated cell death. To further test whether same the UPA-UPA interaction as seen in the rNLRP1-rDPP9 complex is important for downstream NLRP1 inflammasome signaling, we mutated residues of hNLRP1 (hNLRP1^P1278, L1281^) corresponding to those from the UPA dimerization interface of rNLRP1and analyzed the resulting mutants in 293T-ASC-GFP cells using the ASC speck assay described above. As expected, > 30% of the cells formed ASC speck in response to VbP treatment (Fig. 5c and Extended Data Fig. 8b). In contrast, ASC speck formation induced by VbP was substantially reduced in 293T cells expressing either of these two hNLRP1 mutants. Taken together, these data demonstrate that the UPA-UPA dimer interface seen in the 2:1 complex is also required for higher order UPA oligomerization and the activation of the hNLRP1 inflammasome. By extension, these results also suggest that an important way by which DPP9 binding inhibits NLRP1 activation is to ‘capture’ two NLRP1 molecules in a 2:1 complex and prevents it from forming higher order oligomers.

As FIIND protein was monomeric in solution (Extended Data Fig. 7b) in the absence of DPP9, we hypothesized that the ZU5 domain itself contributes directly to inhibiting spontaneous UPA dimerization or oligomerization. Supporting this notion, structural comparison revealed that the two ZU5 domain in the monomeric FIIND conformation (Extended Data Fig. 7b) poses substantial steric hindrance for UPA dimerization as seen in the 2:1 complex (Fig. 5d). In addition, the overlap might also serve to displace the ZU5 subdomain of the 2nd rNLRP1 molecule when docking into the second position (Fig. 2b-d).

## Discussion

Hosts have evolved diverse mechanisms for negative regulation of inflammasome activation to maintain cellular homeostasis^28^. In the current study, we solved the crystal structure of the rNLRP1 FIIND domain as well as the cryo-EM structure of the rNLRP1-rDPP9 complex. These results pinpoint the determinants of rNLRP1-DPP9 interaction and provide a detailed model on how DPP9 interaction maintains NLRP1 in the auto-inhibited state, shedding light on activation of the NLRP1 inflammasome (Extended Data Fig. 8c)

rNLRP1 and rDPP9 form a 2:1 complex in which the FIINDs of two rNLRP1 molecules bind one rDPP9 in distinct conformations. The first rNLRP1 interacts with the DPP9 via the ZU5 domain in the auto-inhibited formation. This allows a second rNLRP1 molecule to complete the complex by interacting with the UPA binding site on rDPP9 and the UPA of the first rNLRP1 molecule. Thus, the 2:1 rNLRP1-rDPP9 complex sequesters the potent UPA-CARD fragments in a dimeric state and prevents them from full-on oligomerization and inflammasome activation (Extended Data Fig. 8c). In fact, sequestration of active domains of NLRs is a general strategy seen in inflammasome regulation^28^. The FIIND of rNLRP1 is monomeric in solution (Extended Data Fig. 7b), suggesting that the FIIND-ZU5 domain has intrinsic auto-inhibitory properties for the prevention of DPP9-indepednent UPA-CARD activation. Further supporting the inhibitory role of the ZU5 domain, structural comparison (Fig. 5d) reveals that ZU5 sterically blocks potential UPA-UPA interactions required for formation of UPA-CARD filaments (Fig. 5b) and NLRP1-mediated cell death (Fig. 5c). Therefore, the ZU5 domain appears to have a critical role in negative regulation of both DPP9-indepdendent and DPP9-depdendent NLRP1 activation, by sterically blocking UPA-mediated formation of functional UPA-CARD filaments and promoting DPP8/9 sequestering of UPA-CARD, respectively. In this scenario, relief of ZU5-mediated inhibition of NLRP1 and CARD8 is expected to be a critical step in their activation.

Taking these data together, we propose that the regulation of NLRP1 abides by the widely accepted ‘guard model’^29^ in plant NLR research, which postulates that NLRs act as ‘guards’ to monitor the integrity of accessory host proteins, or so-called ‘guardees’. In this regard, NLRP1 acts as a guard that constantly monitors the state of its guardee, DPP9. We further postulate that the 2:1 rNLRP1-rDPP9 complex represents the bona fide guard-guardee complex found in mammalian cells. This model predicts that any perturbations of the guardee will trigger the activation of the NLR. This is supported by our functional studies demonstrating that chemical inhibition of DPP9 and disruption of NLRP1-DPP9 binding can both trigger NLRP1 activation. As anthrax LF activates mNlrp1b independently of DPP9, the role of the NLRP1-DPP9 guard-guardee complex in pathogen defense remains to be further elucidated. It is tempting to speculate that a hitherto unknown pathogen effector molecule can trigger NLRP1 activation by inhibiting DPP8/9 enzymatic function and/or disrupting NLRP1-DPP9 binding. If proven, this system would be analogous the regulation of NOD1 or MEFV, which sense and guard the activation state of small Rho GTPases^30,31^. It is also likely that such as an effector molecule might be responsible for the protease-independent activation of rat NLRP1 by *T. gondii*^*17*^.

## Acknowledgments

We thank J. Lei and X. Li at Tsinghua University for data collection. We acknowledge the Tsinghua University Branch of the China National Center for Protein Sciences (Beijing) for providing the cryo-EM facility support. We thank Shanghai Synchrotron Radiation Facility (SSRF) for assistance with data collection. We Thank Meng Han in Technology Center for Protein Science Tsinghua University for Edman sequencing analysis. We also acknowledge generous help and advice from members of Dr. Bruno Reversade’s laboratory at Agency for Science, Technology and Research (A*STAR). This research was funded by the National Natural Science Foundation of China (31421001 to J.C.), the Alexander von Humboldt Foundation (a Humboldt professorship to J.C.), Max Planck-Gesellschaft (a Max Planck fellowship to J.C.), Ministry of Health, Singapore, NMRC grant (MOH-000382-00 to W.B.), Concern Foundation (F.L.Z), Nanyang Assistant Professorship (F.L.Z.) and the National Research Foundation fellowship (F.L.Z.).

## Extended data figures and tables

**Extended data Fig. 1 Complete autocleavage of rNLRP1^FIIND^ in crystals and sequence alignment among rNLRP1, hNLRP1, mNLRP1B and CARD8.**

**a**, Electron density around the active site of rNLRP1^FIIND^. The autocleavage site is between rNLRP1^F968^ and rNLRP1^S969^. **b**, SDS-PAGE analysis of the crystals of rNLRP1^FIIND^. The crystals were harvested and dissolved in SDS-PAGE buffer. Lane 1-4 represent crystals from 4 different wells. **c**, Structural alignment of rNLRP1^FIIND^ and UNC5b. The crystal structure of rNLRP1^FIIND^ was aligned with that of UNC5b (PDB code: 3G5B). Color codes are indicated. **d**, Sequence alignment of the FIIND domains of rNLRP1, hNLRP1, mNLRP1B and CARD8.The rDPP9-interacting residues around the ZU-5 binding site, UPA-binding site and residues at the UPA homodimeric interface are highlighted with red, blue and yellow squares at the bottom, respectively. The two residues from the “FS” catalytic motif are indicated by purple dots at the bottom.

**Extended data Fig. 2 3D reconstruction of the rNLRP1-rDPP9 complex.**

**a**, Full-length GST-rNLRP1 and rDPP9 were expressed in insect cells. The complex was purified through GS4B resin. After elution, GST was removed using precision protease and the complex was subjected to gel filtration. Shown on the left is the gel filtration profile of the complex. The protein fractions were visualized by SDS-PAGE followed by Coomassie-blue staining. **b**, A representative cryo-EM image of the rNLRP1-rDPP9 complex. **c**, Representative views of 2D class averages of the rNLRP1-rDPP9 complex used for 3D reconstruction, 2D classes with different rDPP9 binding features of are shown in different colors. **d**, Flowchart of Cryo-EM data processing and 3D reconstruction of the rNLRP1-rDPP9 complex. **e**, The final EM density map of the rNLRP1-rDPP9 complex. Color coded to show the local resolution estimated by Relion. **f**, FSC curve (at 0.143) of the final reconstruction of the rNLRP1-rDPP9 complex.

**Extended data Fig. 3 3D reconstruction of the rNLRP1^S969A^-rDPP9 complex.**

**a**, A representative cryo-EM image of the rNLRP1^S969A^-rDPP9 complex. **b**, Representative 2D class averages of the rNLRP1^5S969A^-rDPP9 complex. **c**, Flowchart of Cryo-EM data processing and 3D reconstruction of the rNLRP1^S969A^-rDPP9 complex. **d**, The final EM density map of the rNLRP1^S969A^-rDPP9 complex. Color coded to show the local resolution estimated by Relion. **e**, FSC curve (at 0.143) of the final reconstruction of the rNLRP1 ^S969A^-rDPP9 complex.

**Extended data Fig. 4 Structural and sequence alignment.**

**a**, left: Structural alignment between the 1^st^ rDPP9, 2^nd^ rDPP9, hDPP9-APO and hDPP9-1G244 (in cartoon), Color codes are indicated. Better defined loop regions in the rNLRP1-bound rDPP9 subunit are highlighted with open frames. Right: Electron density around the stabilized loop regions induced by NLRP1. **b**, Alignment of the cryo-EM structure of the rNLRP1^FIIND^ (with the ZU5 shown in pink and the UPA in blue) with the crystal structure of the rNLRP1^FIIND^ (in cyan). **c**, Sequence alignment of rDPP9, hDPP9, hDPP8, hDPP4 and hDPP7. The rNLRP1-interacting residues from the ZU5-binding site and UPA-binding site are marked with red and blue squares at the bottom, respectively.

**Extended data Fig. 5 Functional assays to analyze the influence of DPP9 protease activity on DPP9-NLRP1 interaction.**

**a**, Catalytic activity of DPP9 is dispensable for DPP9 binding to NLRP1. The catalytic mutant rDPP9^S729A^ with GST fused at the N-terminus was co-expressed with rNLRP1^FIIND-CARD^ in insect cells. The complex protein was purified through GS4B resin. After elution, GST was removed using precision protease and the complex was subjected to gel filtration. Shown on the left is the gel filtration of the complex. The protein fractions were visualized by SDS-PAGE followed by Coomassie-blue staining. **b**, Mutation of Leu101 of rDPP9 (left) or its equivalent Leu131 of hDPP9 (right) has no effect on their protease activity. Left, 0purified wild-type rDPP9 or the rDPP9^L101E^ mutant protein with concentrations indicated was incubated with a fluorescent test substrate, Gly-Pro-AMC, in buffer containing 25 mM Tris-HCl 8.0, 150 mM NaCl at 25 °C with a reaction volume 100 μl. Samples were taken after 10 min of incubation to measure AMC fluorescence on a spectrometer. On the right, 293T cells were transfected with an empty vector, wild-type hDPP9, hDPP9^S759A^ and hDPP9^L131E^ and lysed in 1xTBS with 0.25% NP-40 48 h after transfection. 0.3μg of total lysate was incubated with a fluorescent test substrate, Gly-Pro-AMC in 50 μL lysis buffer. AMC Fluorescence was continuously measured for 30 mins at room temperature at 1min intervals. The rate of reaction was calculated by the increase of AMC fluorescence per minute (linear fit, R^2^>0.9). **c**, N-terminal sequence analysis of rNLRP1 by Edman degradation. Shown in the left and the right are N-terminal sequences of UPA from rNLRP1^FIIND^ and rNLRP1-rDPP9 determined by Edman degradation, respectively.

**Extended data Fig. 6 Mutagenesis analysis of the ZU5-binding site.**

GST-fused wild type or mutant rNLRP1^FIIND^ and rDPP9 were individually purified from insect cells. Wild type or mutant rNLRP1^FIIND^ protein was used to pull down rDPP9 (including the rDPP9^L101E^ mutant) with GS4B resin. After extensive washing, proteins bound to the GS4B resin were visualized by SDS-PAGE followed by Coomassie-blue staining.

**Extended data Fig. 7 VbP has little effect on rDPP9 interaction with rNLRP1 in vitro, and characterization of rNLRP1^FIIND^ by gel filtration.**

**a**, GST-fused rNLRP1^FIIND^ and rDPP9 were individually purified from insect cells. Wild type or rNLRP1^FIIND^ mutant proteins were used to pull down rDPP9 with GS4B in the presence or absence of VbP. After extensive washing, proteins bound to the GS4B were visualized by SDS-PAGE followed by Coomassie-blue staining. **b**, Gel filtration analyses of the rNLRP1^FIIND^ protein. Shown in left and right panel are gel filtration profile and SDS-PAGE of rNLRP1^FIIND^. Protein fractions visualized by Coomassie-blue staining. Numbers in the left panel indicate the positions of standard protein makers.

**Extended data Fig. 8 293T ASC-GFP specks formation assay and a working model of DPP9-medaited inhibition of NLRP1.**

a, Representative images of DPP8 and DPP9 double knockout (DKO) 293T-ASC-GFP cells transfected with hNLRP1 and wild-type hDPP9 or hDPP9 mutants. Cells were fixed 24 h post transfection. The percentage of cells with ASC-GFP specks were quantified with >200 cells. b, Representative images of 293T-ASC-GFP cells transfected with hNLRP1 mutants indicated. VbP (3 μM) was added 24 h post transfection. Cells were fixed 48 h post transfection and nuclei counterstained with Hoescht 33342. The percentage of cells with ASC-GFP specks were quantified with >200 cells. c, Working model on DPP9-mediated inhibition of NLRP1 and pathogen-induced activation of NLRP1. In resting cells, an autoinhibited rNLRP1 interacts with a dimeric rDPP9 via its autoinhibitory ZU5 domain (A). This interaction enables rDPP9 to recruit an autocleaved rNLRP1, resulting in dissociation of the N-terminal segment including the autoinhibitory ZU5 domain from the C-terminal UPA-CARD and formation of a 2:1 rNLRP1-rDPP9 complex (B). The two active UPA-CARD fragments in the complex are sequestered from oligomerization by interaction with the active sites of rDPP9 and interaction with the DPP9-bound autoinhibited rNLRP1 via UPA-UPA dimerization. As both binding and protease activity are important for rDPP9 to suppress NLRP1, the 2:1 complex is proposed to act as an innate immune surveillance receptor to sense perturbations in NLRP1-DPP9 interaction or DPP9 protease activity by pathogen- or likely host-derived activity. For example, proteasomal degradation of the autoinhibitory ZU5 domain induced by Bacillus anthracis lethal factor (LF) or by the Shigella flexneri-secreted ubiquitin ligase IpaH7.8 would lead to release of the active UPA-CARD fragments from the complex (C). The released UPA-CARD then oligomerize (D) to recruit ASC (E) for activation of downstream immune signaling. Note: for clarity, only one DPP9 monomer is shown to bind two NLRP1 molecules.

## Methods

### Protein expression and purification

The genes of full-length rat NLRP1(rNLRP1) and full-length rat DPP9 (rDPP9) were synthesized by GENEWIZ. The constructs of rNLRP1 (residues 1-1218), rDPP9 (residues 1-862, all mutants), rNLRP1^FIIND^ (residues 822-1122, all mutants) and rNLRP1^FIIND-CARD^ (residues 1822-1218) were generated by standard PCR-based cloning strategy and cloned into pFastBac-1 vector with an N-terminal GST tag or no tag, and their identities were confirmed by sequencing. All the proteins were expressed using the Bac-to-Bac baculovirus expression system (Invitrogen) in sf21 cells at 28 °C. one liter of cells (2.5×10^6^ cells ml^−1^, medium from Expression Systems) was infected with 20 ml baculovirus at 28 ℃. After growth at 28 °C for 48 h, the cells were harvested, re-suspended in the buffer containing 25 mM Tris-HCl pH 8.0 and 150 mM NaCl, and lysed by sonication. The soluble fraction was purified from the cell lysate using Glutathione Sepharose 4B beads (GS4B, GE Healthcare). The proteins were then digested with PreScission protease (GE Healthcare) to remove the GST tag and further purified by gel filtration (SuperoseTM 6 prep grade XK 16/70; GE Healthcare). To prepare for crystallization trials, the purified rNLRP1^FIIND^ (residues 822-1122) protein was concentrated to about 8.0 mg ml-1 in buffer containing 100 mM NaCl, 10 mM Tris-HCl pH 8.0. For co-expression of rNLRP1 and rDPP9, one liter of sf21 cells were co-infected with 10 ml recombinant baculovirus of rNLRP1 and rDPP9, and then rNLRP1-rDPP9 complex was purified using Glutathione Sepharose 4B beads. For cryo-EM investigation, the purified rNLRP1-rDPP9 complex was concentrated to about 0.3 mg/mL in buffer containing 25 mM Tris-HCl pH 8.0, 150 mM NaCl and 3 mM DTT.

Recombinant hNLRP1^UPA-CARD^ tagged with a removable SNAP domain was expressed using bacterial vectors as the form of inclusion bodies. After cellular lysis, the cellular pellet was harvested after 30,000 g centrifugation for 30 min at 4 degree. Additional wash using wash buffer (20 mM Tris-HCl, pH 8.0, 150 mM NaCl, 1% Triton-X and 1 mM DTT) several times until pure white pellet was obtained. The pellet was dissolved in 6 M guanidinium, and centrifugated at 30,000 g for a 2^nd^ time for 30 min at room temperature to remove contaminants. The denatured soluble proteins were then gradually dialyzed against 3, 2, 1.5, 1, 0.8, and 0.6 M guanidinium containing dialysis buffer (20 mM Tris-HCl, pH 8.0, 150 mM NaCl, 5 mM beta-mercaptoethanol) in the cold room, and eventually in fresh dialysis buffer without granidinium. The refolded proteins were centrifugated for the third time at 10,000g for 10 min to get rid of mis-folded aggregates. The soluble refolded fractions were then subjected to biochemical analysis and negative stain EM experiments. SNAP tag was removed by 3C proteases, and further purified away by reverse Ni-NTA purification.

### Gel filtration assay

The GST-rNLRP1^FIIND S969A^-rDPP9 complex and rNLRP1^FIIND^ proteins purified as described above were subjected to gel filtration (Superose 6, 10/30; GE Healthcare) in buffer containing 10 mM Tris pH 8.0 and 100 mM NaCl. The purified rNLRP1^FIIND^ protein was left at 18 °C for two weeks to obtain its fully autoprcoessed form. The fully processed rNLRP1^FIIND^ was then incubated with the purified GST-rNLRP1^FIIND S969A^-rDPP9 complex with a molar ratio of about 1:1 in 4 °C for 150 min before gel filtration analysis. Samples from relevant fractions were applied to SDS PAGE and visualized by Coomassie blue staining. A similar procedure was used to verify the interaction of rDPP9 with other rNLRP1 mutant proteins.

### Pull-down assay

50 ml of sf21 cells (2.5×10^6^ cells ml^−1^, medium from Expression Systems) were infected with 1 ml baculovirus of GST-rNLRP1^FIIND^ wild-type or mutants, and then were expressed and purified as described above. The proteins were purified from the cell lysate using 300 μl GST resin (GS4B, GE Healthcare), and incubated with excess purified wild-type or mutant rDPP9 proteins on ice for 60 min. The resins were washed with 1 ml buffer containing 10 mM Tris pH 8.0, 100 mM NaCl for five times, and were eluted with 300 μl buffer containing 25 mM Tris pH 8.0, 150 mM NaCl, 15 mM GSH. The eluted samples were analyzed by SDS-PAGE and visualized by Coomassie blue staining.

To test effect of VbP on rNLRP1-rDPP9 interaction, 2 mM VbP was added to the purified rDPP9, GST-rNLRP1^FIIND S969A^-rDPP9 complex or GST-rNLRP1^FIIND^-rDPP9 complex. After 60 min incubation, the samples were individually incubated with wild type or mutant rNLRP1^FIIND^ protein and 100 μl GS4B resin on ice for 60 min. After extensive washing, the proteins bound in the resin were eluted and analyzed by SDS-PAGE and visualized by Coomassie blue staining.

### Enzymatic activity assay

To measure rDPP9 activity, stock solution of substrate (10 mM Gly-Pro-AMC) was prepared in DMSO. The purified wild-type or mutant rDPP9 protein was diluted to 0.5 μM, 1 μM and 5 μM into a final volume of 100 μl in buffer containing 10 mM Tris pH 8.0 and 100 mM NaCl. 10 μl of 10 μM substrate Gly-Pro-AMC dissolved in DMSO was added to the mixture. Substrate cleavage was measured as liberated AMC fluorescence signal recorded at room temperature in a Luminescence Spectrometer at 380-nm excitation and 500-nm emission wavelengths over a 10 min.

To measure the activity of hDPP9, 293T cells were transfected with hDPP9 and lysed 48 hours post transfection in 1x Tris-buffered saline (TBS) with 0.25% NP40. 0.3μg of lysate was mixed with 0.1 μl 100mM Gly-Pro-AMC in 50 μL lysis buffer. AMC fluorescence (380 nm excitation; 460 nm emission) was monitored at room temperature for 30 mins at 1 min intervals.

### Edman degradation by the PPSQ-33A system

The phenylthiohydantoin amino acid is separated in the reversed-phase mode of high-performance liquid chromatography using the differences between the retention times of different amino acids, and the amount of UV absorbance for specific wavelengths was detected. The samples were transferred to PVDF membrane and 5 cycles was set. The amino acid sequences of each sample were determined from the chromatograms obtained in each cycle evaluation performed by comparing chromatograms with those in previous and subsequent cycles and identifying the PTH-amino acids with the greatest increase.

### Cryo-EM sample preparation and data collection

An aliquot of 3 μl purified protein of rNLRP1-rDPP9 or rNLRP1^FIIND S969A^-rDPP9 complex was applied to holey carbon grids (Quantifoil Au 1.2/1.3, 300 mesh), which were glow-discharged for 30 s at middle level in Harrick Plasma after 2 min evacuation. The grids were then blotted by filter papers (TED PELLA, INC.) for 2.5 s in 8 °C with 100% humidity and flash-frozen in liquid ethane using FEI Vitrobot Marked IV.

Cryo-EM data for rNLRP1-rDPP9 and rNLRP1^FIIND S969A^-rDPP9 complex was collected on Titan Krios electron microscope operated at 300 kV, equipped with a Gatan K2 Summit direct electron detector and a Gatan Quantum energy filter (an additional Cs-corrector for rNLRP1^FIIND S969A^-rDPP9 data collection). A total of 7,157 and 4,971 micrograph stacks were automatically recorded using AutoEMation in super-resolution mode for rNLRP1-rDPP9 and rNLRP1^FIIND S969A^-rDPP9, at a nominal magnification of 130,000 × and 105,000 ×, respectively. Defocus values varied from −1.0 μm to −2.0 μm for both data set^32^. Dose rates for rNLRP1-rDPP9 and rNLRP1^FIIND S969A^-rDPP9 data collection were 10.008 and 10.621 electron per pixel per second, respectively. For both data sets the exposure time of 5.6 s was dose-fractionated into 32 sub-frames, leading to a total accumulated dose of approximate 50 electrons per Å^2^ for each stack.

### Image processing and 3D reconstruction

The stacks of rNLRP1-rDPP9 and rNLRP1^FIIND S969A^-rDPP9 recorded in super-resolution mode were motion-corrected by MotionCor2 and binned twofold, resulting in a physical pixel size of 1. 061 Å per pixel and 1.091 Å per pixel, respectively^33^. Meanwhile dose weighting for the summed micrographs was performed^34^. CTFFIND4 was then used to estimate the contrast transfer function (CTF) parameters^35^. Based on CTF estimation, 7,033 and 4,667 micrographs were manually selected for rNLRP1-rDPP9 and rNLRP1^FIIND S969A^-rDPP9, respectively, and were further processed in RELION3.1. Approximately 2,000 particles were manually picked and 2D-classified to generate initial templates for auto-picking. In the end, 2,700,586 and 1,725,380 particles were automatically picked for rNLRP1-rDPP9 and rNLRP1^FIIND S969A^-rDPP9, respectively, using RELION. After several rounds of reference-free 2D classification, 1,430,734 particles for rNLRP1-rDPP9 and 1,117,656 particles for rNLRP1^FIIND S969A^-rDPP9 were subjected to 3D classification, using initial 3D reference models obtained by ab initio calculation from RELION3.1. Particles from good 3D classes with better overall structure features were selected for 3D refinement. After global 3D refinement and post-processing, the resolution was 3.07 Å with a particle number of 343,648 for rNLRP1-rDPP9, and 3.69 Å with a particle number of 252,425 for rNLRP1^FIIND S969A^-rDPP9.

To improve the density quality of the NLRP1 part in the rNLRP1-rDPP9 map, the rNLRP1-rDPP9 particles after 3D refinement was then subjected to a further round of focused 3D classification with a local mask generated using Chimera, the procedure of the focused 3D classification previously reported was adopted to select the 3D class with good density^36^. Ultimately, a subset of 182,116 particles after focused 3D classification were subjected to a final 3D refinement and yielded a global reconstruction at 3.18 Å after postprocess.

2D classification, 3D classification and 3D auto-refinement were all performed with RELION3.1(Ref. 37-39). The resolutions were determined by gold-standard Fourier shell correlation^40^. Local resolution distribution was evaluated using RELION^41^.

### Crystallization, data collection and structure determination

Crystallization of rNLRP1 was performed by hanging-drop vapour-diffusion methods using mixing 1 μl of 8 mg/ml protein with 1 μl of reservoir solution at 18 °C. Good quality crystals of rNLRP1^FIIND^ were obtained in buffer containing 1.0 M Ammonium sulfate, 0.1 M Bis-Tris pH 5.5, 1% w/v polyethylene glycol 3,350. All the crystals were flash frozen in the condition of the 15% glycerol added reservoir buffer as the cryo-protectant to prevent radiation damage. The diffraction data set was collected at Shanghai Synchrontron Radiation Facility (SSRF) on the beam line BL19U1 using a CCD detector and was processed using HKL2000 software package. The crystal structure of rNLRP1^FIIND^ was determined by PHASER_MR with the structure of NUC5b^18^ as the search model. The model from the MR was manually rebuilt to the sequence of rNLRP1^FIIND^ in the program COOT^42^ and subsequently subjected to refinement by the program Refine_Phenix^43^. Data collection, processing, and refinement statistics are summarized in Extended Data Table 1.

### Model building and refinement

The EM density map of rNLRP1-rDPP9 was used for model building as its quality of the EM density of rNLRP1 was good enough for sequence assignment. The model of hDPP9 (PDB:6EOQ)^22^ along with two copies of rNLRP1^FIIND^ crystal structure we determined before were docked into the EM density map of rNLRP1-rDPP9 in Chimera^44^. The sequence of hDPP9 was changed to that of rDPP9, the whole model containing two rNLRP1^FIIND^ molecules and a rDPP9 dimer was then adjusted manually in the program COOT^42^, and refined against the EM map by PHENIX in real space with secondary structure and geometry restraints^43^. The final model of the rNLRP1-rDPP9 complex was validated using MolPorbity and EMRinger in PHENIX package^43^. Extended Data Table 2 summarizes the model statistics.

### 293T-ASC-GFP transfection and ASC-GFP speck formation assay

293T-ASC-GFP or DPP8 DPP9 DKO cells were transfected with the indicated plasmids with Lipofectamine 2000 (ThermoFisher). For immunoprecipitation, cells were harvested 48 h post transfection. For the ASC-GFP speck assay, cells were fixed 24 h post transfection and counterstained with DAPI or Hoescht before wide field fluorescence imaging. The number of nuclei per field of view was counted in ImageJ via the following image processing steps: 1) Threshold (20~30 to 255), 2) Watershed and 3) Analyze Particles (200-Infinity). ASC speck were counted in ImageJ in the GFP channel using Find Maxima (Prominence=20).

### Data availability

The cryo-EM density map of the wild type rNLRP1-rDPP9 complex and its corresponding coordinates have been deposited in the Electron Microscopy Data Bank (EMDB) and the Protein Data Bank (PDB), respectively, with the following accession codes: EMD-30458 and 7CRW. The coordinates of the crystal structure of the FIIND of rNLRP1 has been deposited in the PDB with the accession code 7CRV. The cryo-EM density map of the FIIND-CARD S969A mutant fragment bound by rDPP9 has been deposited in EMDB with the accession code EMD-30459.

