## Supplementary figures and images for "Structural and biochemical mechanisms of NLRP1 inhibition by DPP9"

### Extended Data Fig 1

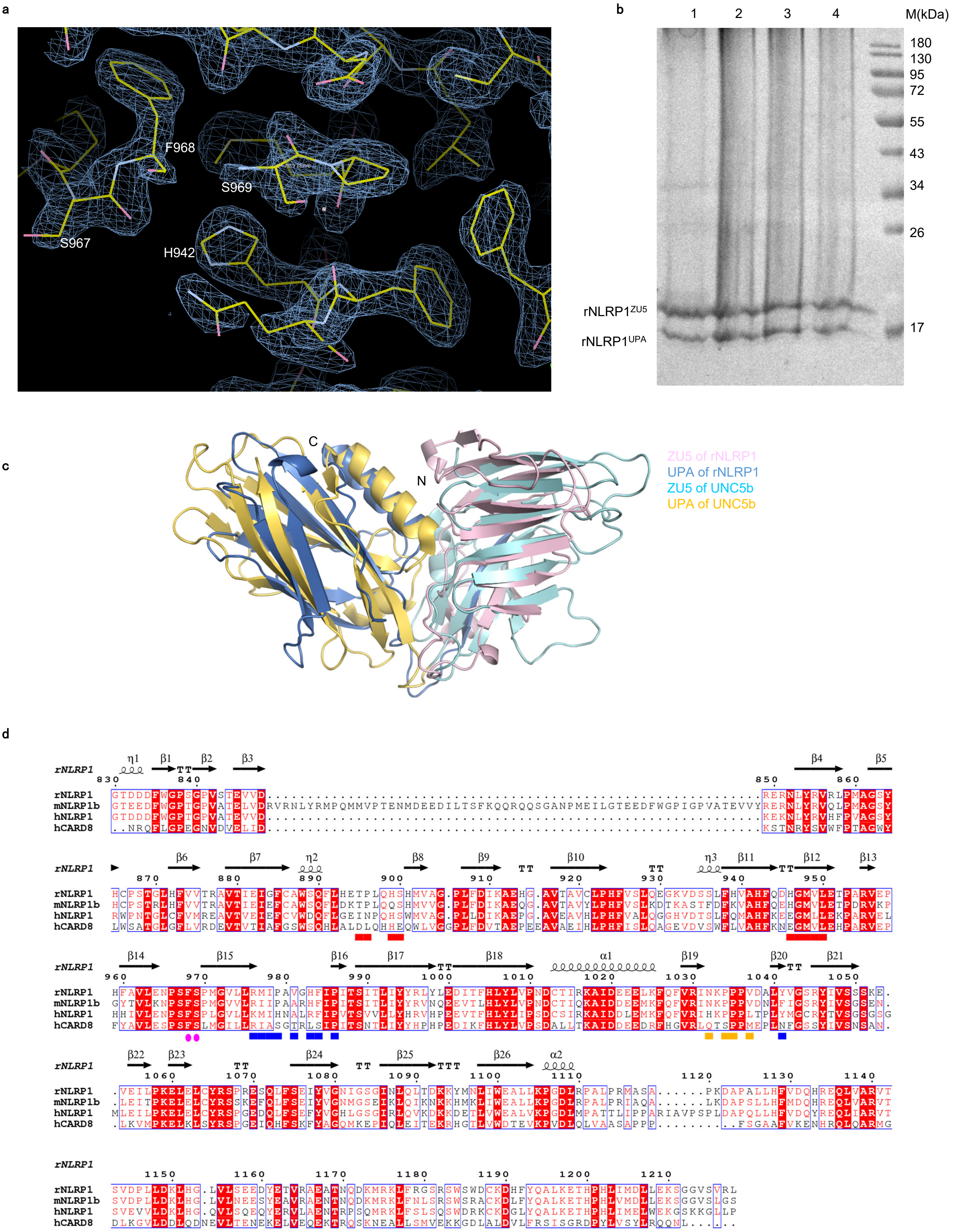

### Extended Data Fig 2

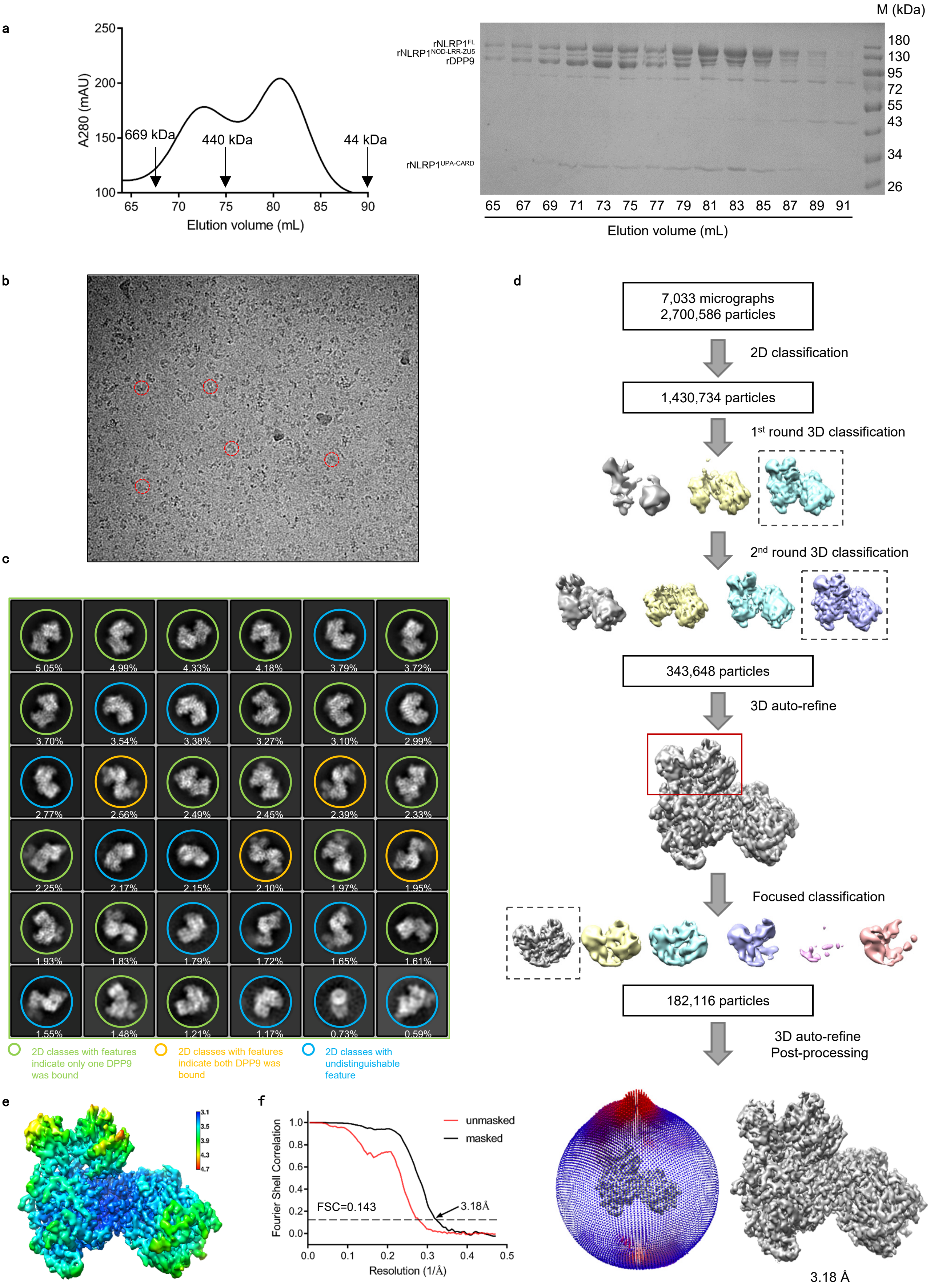

### Extended Data Fig 3

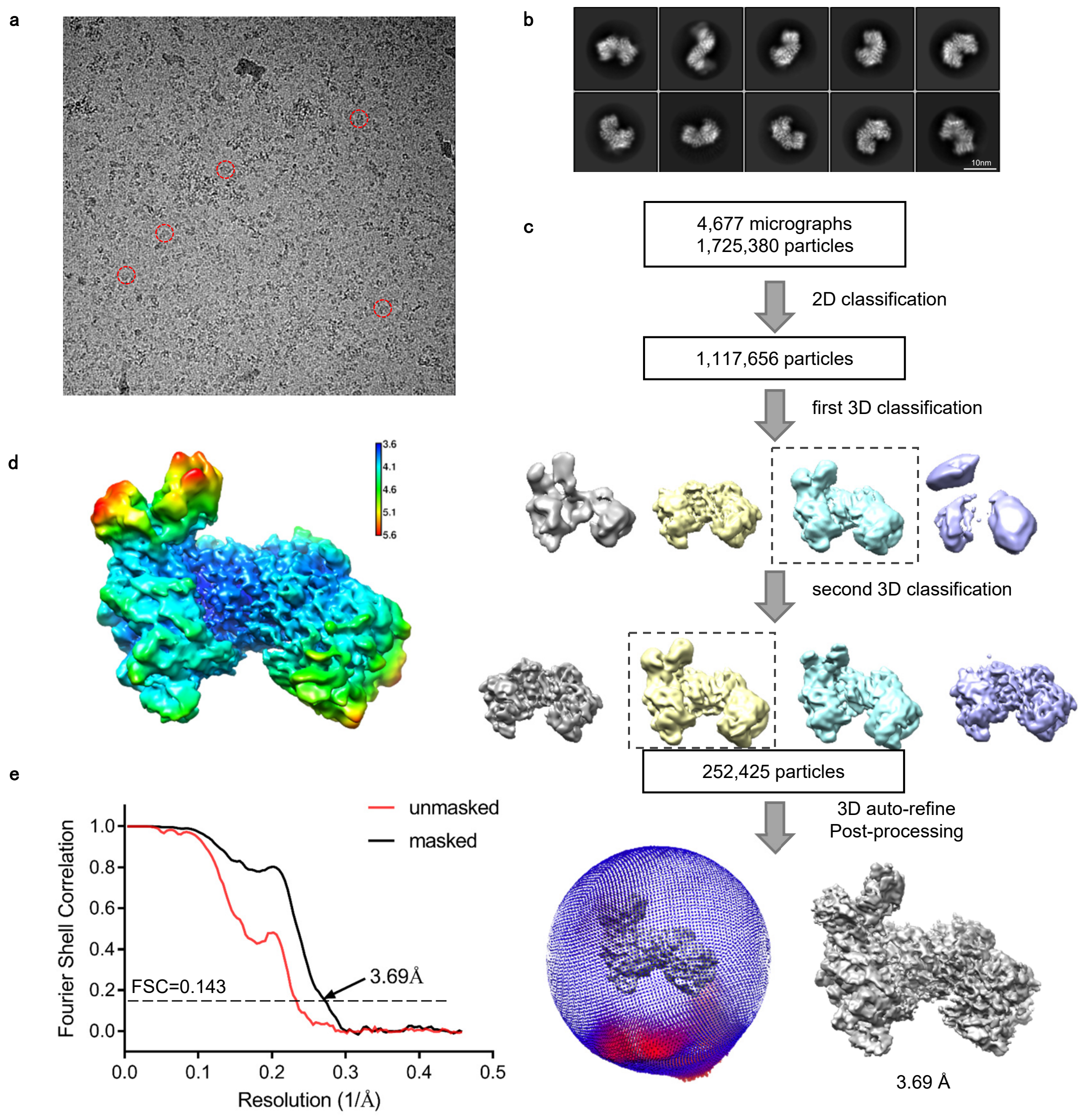

### Extended Data Fig 4

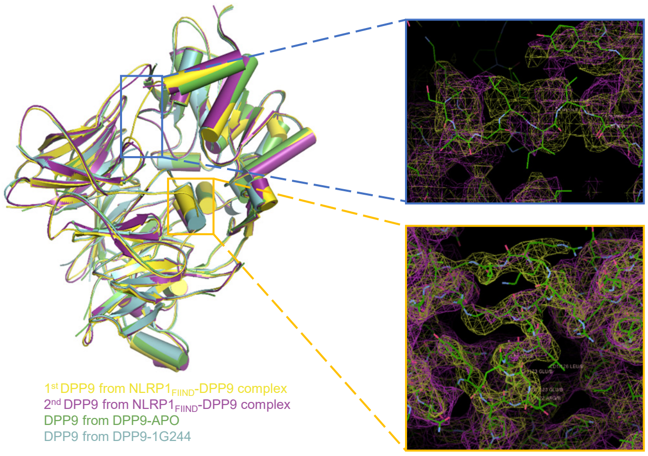

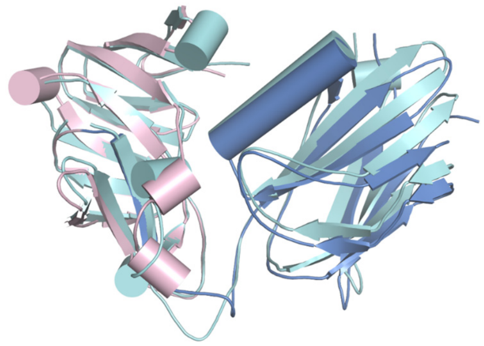

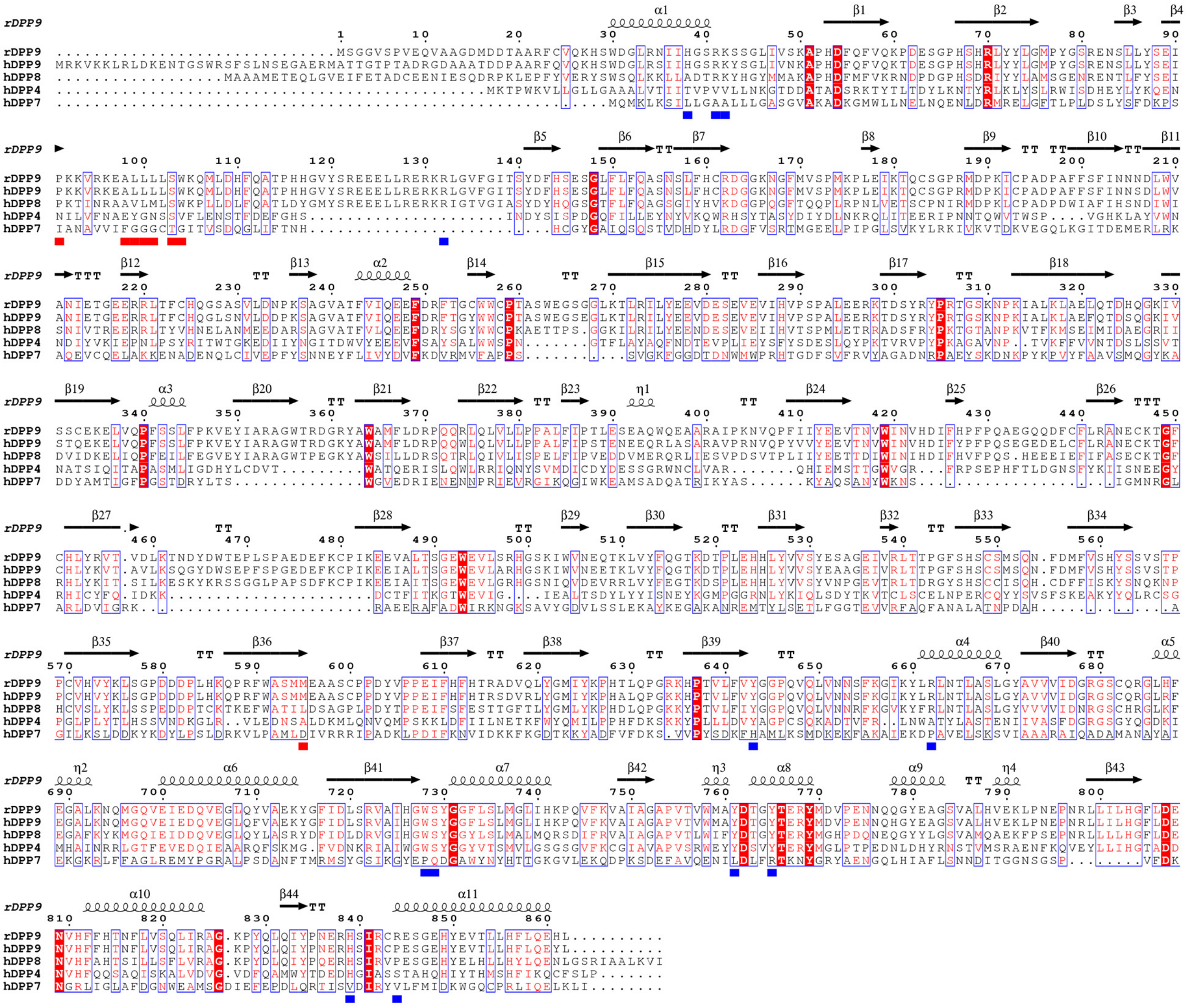

### Extended Data Fig 6

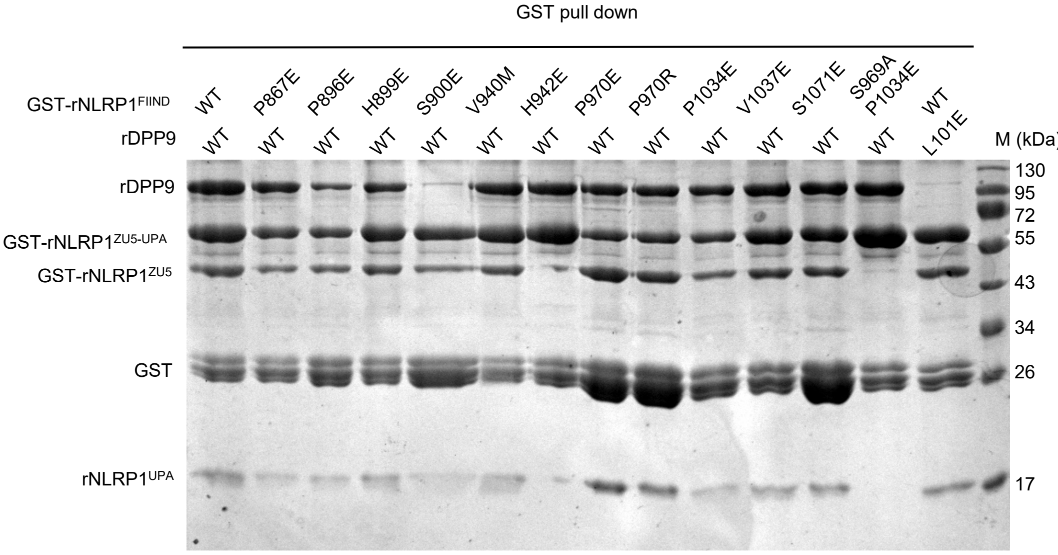

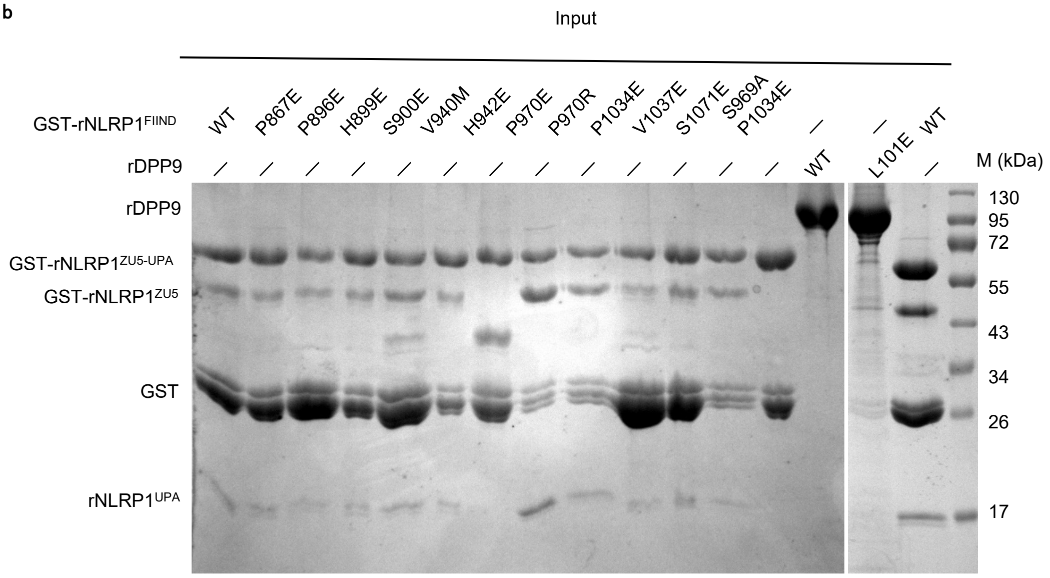

### Extended Data Fig 7

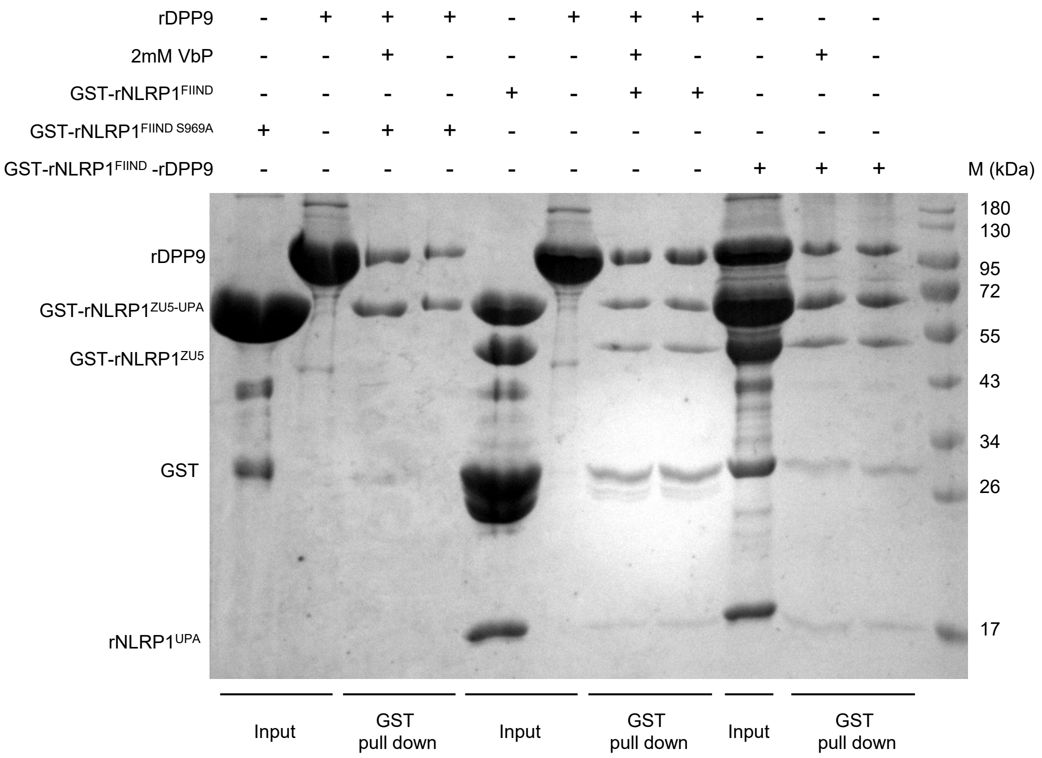

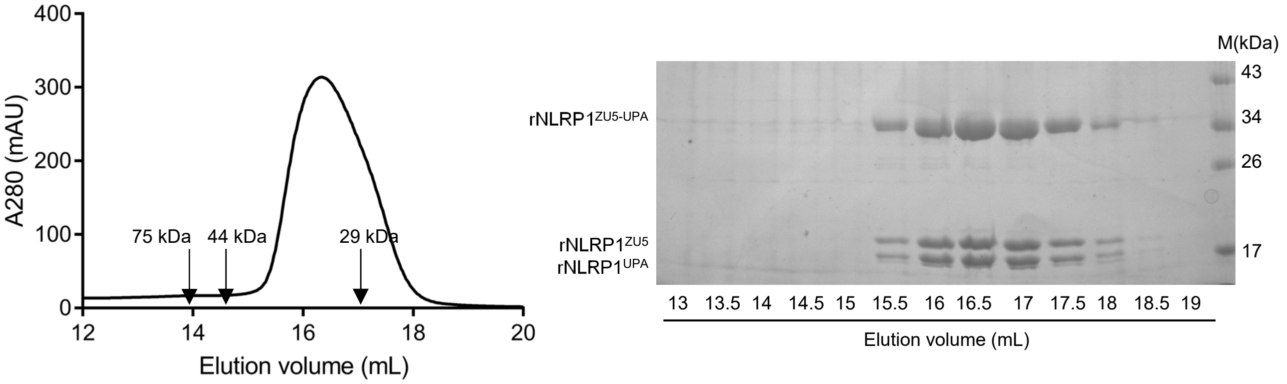

### Extended Data Fig 8

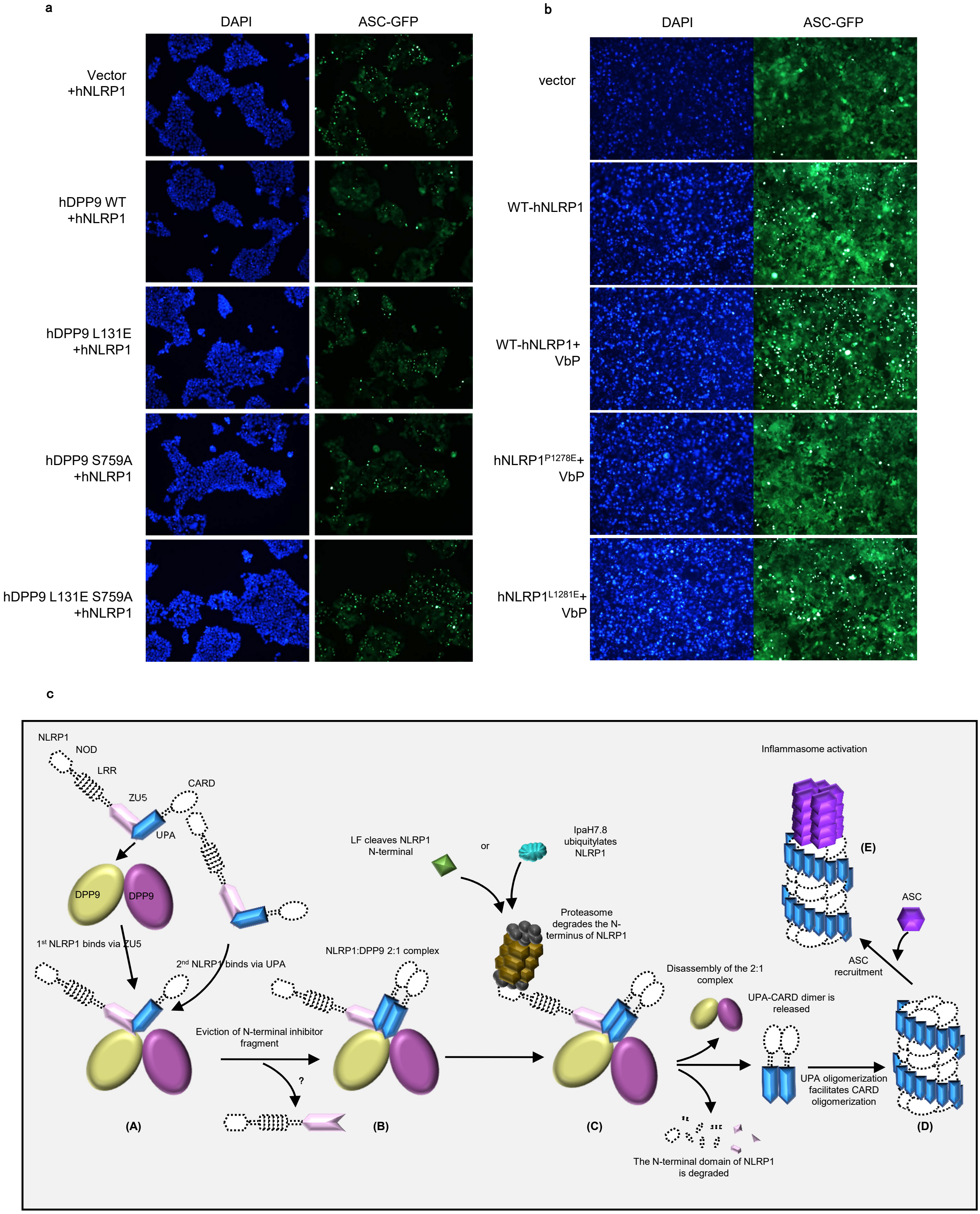
