## Extended Data Fig 5 for "Structural and biochemical mechanisms of NLRP1 inhibition by DPP9"

a

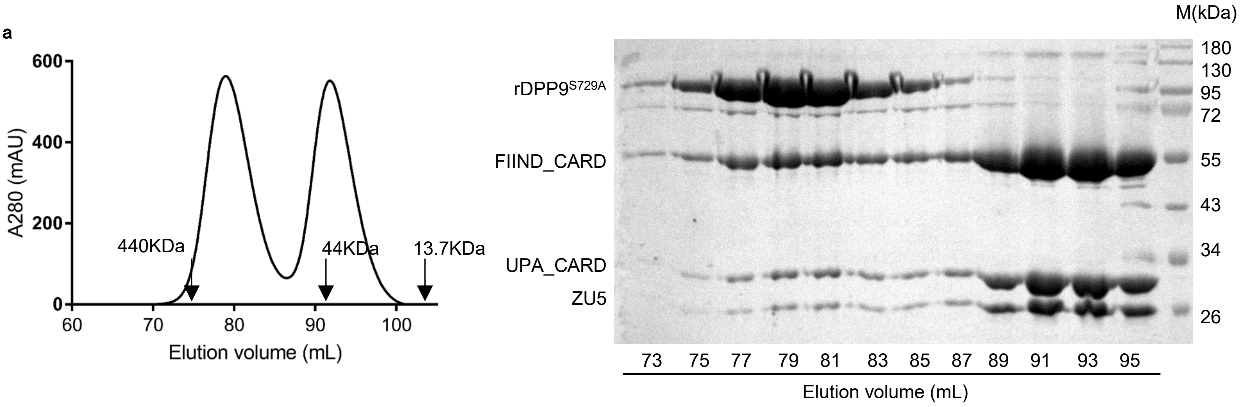


b

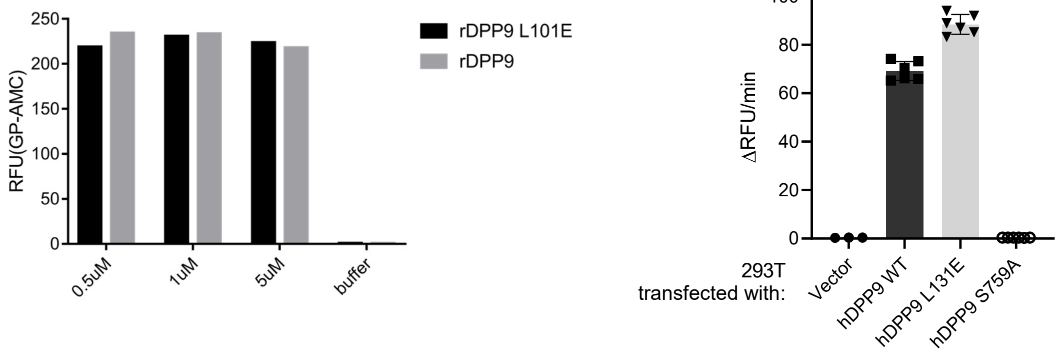


c

rNLRP1$_{FIIND}$ monomer

[Sequence]
Ser Pro Met Gly Val

[Estimated Sequence]

| | 1 | 2 | 3 | 4 | 5 |
|---|---|---|---|---|---|
| 1st | Ser | Pro | Met | Gly | Val |
| 2nd | Gly | Asp | Thr | Tyr | Leu |
| 3rd | Asp | Phe | Ile | Asn | Ala |
| 4th | His | Glu | Leu | Glu | Tyr |
| Reliability(%) | 20.9 | 100.0 | 79.0 | 72.4 | 100.0 |

[Evaluated Value]

| | 1 | 2 | 3 | 4 | 5 |
|---|---|---|---|---|---|
| Asp | 5.52 | 3.19 | 0.64 | 0.70 | 1.12 |
| Glu | 0.73 | 1.08 | 1.04 | 1.12 | 1.11 |
| Asn | 0.24 | 0.52 | 0.67 | 1.20 | 0.78 |
| Gln | 3.88 | 0.44 | 1.39 | 0.63 | 0.96 |
| Ser | 18.30 | 0.11 | 0.18 | 0.54 | 1.00 |
| Thr | 2.24 | 0.47 | 2.24 | 0.41 | 1.25 |
| His | 4.59 | 0.23 | 0.92 | 0.00 | 0.00 |
| Gly | 7.23 | 0.51 | 0.74 | 11.65 | 0.44 |
| Ala | 2.61 | 0.29 | 1.42 | 0.38 | 1.43 |
| Tyr | 0.87 | 0.86 | 1.04 | 1.40 | 1.26 |
| Arg | 1.29 | 0.20 | 1.23 | 0.62 | 0.83 |
| Met | 1.25 | 0.88 | 19.94 | 0.21 | 0.18 |
| Val | 0.62 | 0.98 | 1.15 | 0.84 | 41.15 |
| Pro | 0.03 | 47.23 | 0.32 | 0.27 | 0.47 |
| Trp | 1.08 | 0.45 | 0.00 | 0.00 | 0.00 |
| Phe | 0.83 | 1.19 | 1.40 | 0.62 | 0.91 |
| Lys | 1.05 | 0.73 | 1.37 | 0.18 | 0.39 |
| Ile | 0.95 | 1.06 | 2.15 | 0.83 | 0.94 |
| Leu | 0.98 | 1.00 | 1.74 | 0.58 | 2.33 |

rNLRP1$_{FIIND}$-rDPP9 complex

[Sequence]
Ser Pro Met Gly Val

[Estimated Sequence]

| | 1 | 2 | 3 | 4 | 5 |
|---|---|---|---|---|---|
| 1st | Ser | Pro | Met | Gly | Val |
| 2nd | Leu | Ile | Ile | Asn | Leu |
| 3rd | Gln | Ala | Tyr | Ile | Ala |
| 4th | Gly | Glu | Glu | Glu | Tyr |
| Reliability(%) | 36.9 | 68.6 | 81.2 | 48.8 | 100.0 |

[Evaluated Value]

| | 1 | 2 | 3 | 4 | 5 |
|---|---|---|---|---|---|
| Asp | 0.20 | 1.93 | 0.58 | 0.78 | 0.84 |
| Glu | 0.69 | 2.33 | 1.77 | 1.32 | 0.80 |
| Asn | 0.05 | 0.48 | 0.89 | 2.78 | 0.55 |
| Gln | 3.18 | 0.50 | 0.88 | 0.78 | 0.86 |
| Ser | 19.37 | 0.19 | 0.31 | 0.83 | 0.90 |
| Thr | 1.09 | 0.92 | 1.65 | 1.07 | 0.73 |
| His | 2.49 | 0.14 | 0.84 | 0.34 | 1.01 |
| Gly | 2.87 | 0.88 | 0.60 | 13.75 | 0.38 |
| Ala | 0.64 | 2.49 | 0.73 | 0.77 | 2.10 |
| Tyr | 0.93 | 1.09 | 2.37 | 1.03 | 1.87 |
| Arg | 0.42 | 2.09 | 0.88 | 0.69 | 0.73 |
| Met | 1.01 | 0.99 | 25.91 | 0.31 | 0.02 |
| Val | 0.86 | 1.22 | 1.41 | 1.26 | 37.73 |
| Pro | 0.06 | 26.14 | 0.40 | 0.45 | 1.33 |
| Trp | 1.82 | 0.60 | 0.38 | 0.00 | 0.00 |
| Phe | 0.57 | 1.84 | 0.68 | 1.27 | 1.00 |
| Lys | 1.14 | 0.17 | 1.18 | 0.25 | 1.09 |
| Ile | 0.43 | 3.37 | 3.41 | 2.13 | 0.58 |
| Leu | 4.71 | 0.40 | 1.00 | 1.13 | 2.40 |
