## Extended Data table 1 for "Structural and biochemical mechanisms of NLRP1 inhibition by DPP9"

Extended Data Table 1: X-ray diffraction data collection and refinement statistics

| | Crystal for NLRP1_FIIND |
|---|---|
| PDB ID | 7CRV |
| **Data collection** | |
| Space group | P3121 |
| Cell dimensions | |
| *a, b, c* (Å) | 83.96,83.96,156.54 |
| *α*, *β*, *γ* (°) | 90,90,120 |
| Resolution range(Å) | 50.00-2.00 (2.03-2.00) |
| $R_{sym}$ (%) | 8.7 (99) |
| I/δI | 30.9 (2.2) |
| Completeness (%) | 99.9 (96.5) |
| Redundancy | 19.3 (16.8) |
| **Refinement** | |
| Resolution (Å) | 28.76-2.00 (2.07-2.00) |
| No. reflections | 43788 (4250) |
| $R_{work}/R_{free}$ (%) | 22.45/25.91 (29.44/34.86) |
| No. atoms | 4542 |
| Protein residues | 571 |
| B-factors | 37.34 |
| R.M.S deviations | |
| Bonds lengths (Å) | 0.011 |
| Bonds angles (°) | 1.40 |
| Ramachandran plot statistics | |
| Preferred (%) | 97.87 |
| Allowed (%) | 1.95 |
| Outlier (%) | 0.18 |
