## Extended Data table 2 for "Structural and biochemical mechanisms of NLRP1 inhibition by DPP9"

Extended Data Table 2: Cryo-EM data collection, 3D reconstruction and model statistics

| | Nlrp1-DPP9 | NLRP1-FIIND(S969A)-DPP9 |
|---|---|---|
| PDB ID | 7CRW | NA |
| EMDB ID | EMD-30458 | EMD-30459 |
| **Data collection** | | |
| Cryo electron microscope | FEI Titan Krios | FEI Titan Krios |
| Voltage (kV) | 300 | 300 |
| Detector | Gatan K2 Summit with Gatan GIF Quantum (20eV slit) | Gatan K2 Summit with Gatan GIF Quantum (20eV slit) and Cs corrector |
| Magnification | 130,000 | 105,000 |
| Pixel size (Å) | 1.061 | 1.091 |
| Total electron dose ($e^-$/Å$^2$) | 49.784 | 49.969 |
| Exposure rate ($e^-$/pixel/sec) | 10.008 | 10.621 |
| Defocus range (μm) | -1.0 ~ -2.0 | -1.0 ~ -2.0 |
| Micrographs collected | 7,157 | 4,971 |
| Micrographs used | 7,033 | 4,667 |
| **3D reconstruction** | | |
| Software | RELION 3.1 | RELION 3.1 |
| Total extraced particles | 2,700,586 | 1,725,380 |
| Number of particles used for 3D reconstruction | 182,116 | 252,425 |
| Symmetry imposed | C1 | C1 |
| Resolution range (Å) | 3.06-4.65 | 3.60-6.14 |
| Resolution (Å) after refinement | 3.64 (FSC=0.143) | 4.29(FSC=0.143) |
| Resolution (Å) after post-processing | 3.18 (FSC=0.143) | 3.69(FSC=0.143) |
| Map sharpening B-factor (Å$^2$) | -50 | -100 |
| **Refinement and validation** | | |
| Software | Phenix.real_space_refine | NA |
| Model resolution (Å) | 3.3 (FSC=0.5) | NA |
| Model composition | | |
| Non-hydrogen atoms | 16,614 | NA |
| Protein residues | 2057 | NA |
| Map-model CC (overall/local) | 0.85/0.85 | NA |
| Rwork/Rfree (%) | 33.41/33.41 | NA |
| B factors | 80.08 | NA |
| R.M.S deviations | | |
| Bonds lengths (Å) | 0.005 | NA |
| Bonds angles (°) | 0.723 | NA |
| MolProbity score | 1.90 | NA |
| Clash score | 9.23 | NA |
| Rotamer outliers (%) | 0.60 | NA |
| EMRinger score | 3.00 | NA |
| Ramachandran plot statistics | | |
| Preferred (%) | 93.91 | NA |
| Allowed (%) | 5.94 | NA |
| Outlier (%) | 0.15 | NA |
